# Distribution and diversity of dimetal-carboxylate halogenases in cyanobacteria

**DOI:** 10.1101/2021.01.05.425448

**Authors:** Nadia Eusebio, Adriana Rego, Nathaniel R. Glasser, Raquel Castelo-Branco, Emily P. Balskus, Pedro N. Leão

## Abstract

Halogenation is a recurring feature in natural products, especially those from marine organisms. The selectivity with which halogenating enzymes act on their substrates renders halogenases interesting targets for biocatalyst development. Recently, CylC – the first predicted dimetal-carboxylate halogenase to be characterized – was shown to regio- and stereoselectively install a chlorine atom onto an unactivated carbon center during cylindrocyclophane biosynthesis. Homologs of CylC are also found in other characterized cyanobacterial secondary metabolite biosynthetic gene clusters. Due to its novelty in biological catalysis, selectivity and ability to perform C-H activation, this halogenase class is of considerable fundamental and applied interest. However, little is known regarding the diversity and distribution of these enzymes in bacteria. In this study, we used both genome mining and PCR-based screening to explore the genetic diversity and distribution of CylC homologs. While we found non-cyanobacterial homologs of these enzymes to be rare, we identified a large number of genes encoding CylC-like enzymes in publicly available cyanobacterial genomes and in our in-house culture collection of cyanobacteria. Genes encoding CylC homologs are widely distributed throughout the cyanobacterial tree of life, within biosynthetic gene clusters of distinct architectures. Their genomic contexts feature a variety of biosynthetic partners, including fatty-acid activation enzymes, type I or type III polyketide synthases, dialkylresorcinol-generating enzymes, monooxygenases or Rieske proteins. Our study also reveals that dimetal-carboxylate halogenases are among the most abundant types of halogenating enzymes in the phylum Cyanobacteria. This work will help to guide the search for new halogenating biocatalysts and natural product scaffolds.

**Data statement:** All supporting data and methods have been provided within the article or through a Supplementary Material file, which includes 14 supplementary figures and 4 supplementary tables.

## Introduction

Nature is a rich source of new compounds that fuel innovation in the pharmaceutical and agriculture sectors [1]. The remarkable diversity of natural products (NPs) results from a similarly diverse pool of biosynthetic enzymes [2]. These often are highly selective and efficient, carrying out demanding reactions in aqueous media, and therefore are interesting starting points for the development of industrially-relevant biocatalysts [2]. Faster and more accessible DNA sequencing technologies have enabled, in the past decade, a large number of genomics and metagenomics projects focused on the microbial world [3]. The resulting sequence data holds immense opportunities for the discovery of new microbial enzymes and their associated NPs [4].

Halogenation is a widely used and well-established reaction in synthetic and industrial chemistry [5], which can have significant consequences for the bioactivity, bioavailability and metabolic activity of a compound [5-7]. Halogenating biocatalysts are thus highly desirable for biotechnological purposes [6, 8]. The mechanistic aspects of biological halogenation can also inspire the development of organometallic catalysts [9]. Nature has evolved multiple strategies to incorporate halogen atoms into small molecules [6], as illustrated by the structural diversity of thousands of currently known halogenated NPs, which include drugs and agrochemicals [10, 11]. Until the early 1990’s, haloperoxidases were the only known halogenating enzymes. Research on the biosynthesis of halogenated metabolites eventually revealed a more diverse range of halogenases with different mechanisms. Currently, biological halogenation is known to proceed by distinct electrophilic, nucleophilic or radical mechanisms [6]. Electrophilic halogenation is characteristic of the flavin-dependent halogenases and the heme- and vanadium-dependent haloperoxidases, which catalyze the installation of C-I, C-Br or C-Cl bonds onto electron-rich substrates. Two families of nucleophilic halogenases are known, the halide methyltransferases and SAM halogenases. Both utilize *S*-adenosylmethionine (SAM) as an electrophilic co-factor or as a co-substrate and halide anions as nucleophiles. Notably, these are the only halogenases capable of generating C-F bonds. Finally, radical halogenation has only been described for nonheme-iron/2-oxo-glutarate (2OG)-dependent enzymes. This type of halogenation allows the selective insertion of a halogen into a non-activated, aliphatic C-H bond. A recent review by Agarwal et al (2017) thoroughly covers the topic of enzymatic halogenation.

Cyanobacteria are a rich source of halogenases among bacteria, in particular for nonheme iron/2OG-dependent and flavin-dependent halogenases (Fig. 1). AmbO5 and WelO5 are cyanobacterial enzymes that belong to the nonheme iron/2OG-dependent halogenase family [12-14]. AmbO5 is an aliphatic halogenase capable of site-selectively modifying ambiguine, fischerindole and hapalindole alkaloids [12, 13]. The close homolog (79% sequence identity) WelO5 is capable of performing analogous halogenations in hapalindole-type alkaloids and it is involved in the biosynthesis of welwintindolinone [13, 15]. BarB1 and BarB2 are also nonheme iron/2OG-dependent halogenases that catalyze trichlorination of a methyl group from a leucine substrate attached to the peptidyl carrier protein BarA in the biosynthesis of barbamide [16-18]. Other halogenases from this enzyme family include JamE, CurA, and HctB. JamE and CurA catalyse halogenations in intermediate steps of the biosynthesis of jamaicamide and curacin A, respectively [19, 20], while HctB is a fatty acid halogenase responsible for chlorination in hectochlorin assembly [21]. ApdC and McnD are FAD-dependent halogenases responsible for the modification of cyanopeptolin-type peptides (also known as (3*S*)-amino-(6*R*)-hydroxy piperidone (Ahp)-cyclodepsipeptides). These enzymes halogenate, respectively, anabaenopeptilides in *Anabaena* and micropeptins in *Microcystis* strains [22-25]. AerJ is another example of a FAD-dependent halogenase, which acts during aeruginosin biosynthesis in *Planktothrix* and *Microcystis* strains [24].

**Figure 1.**
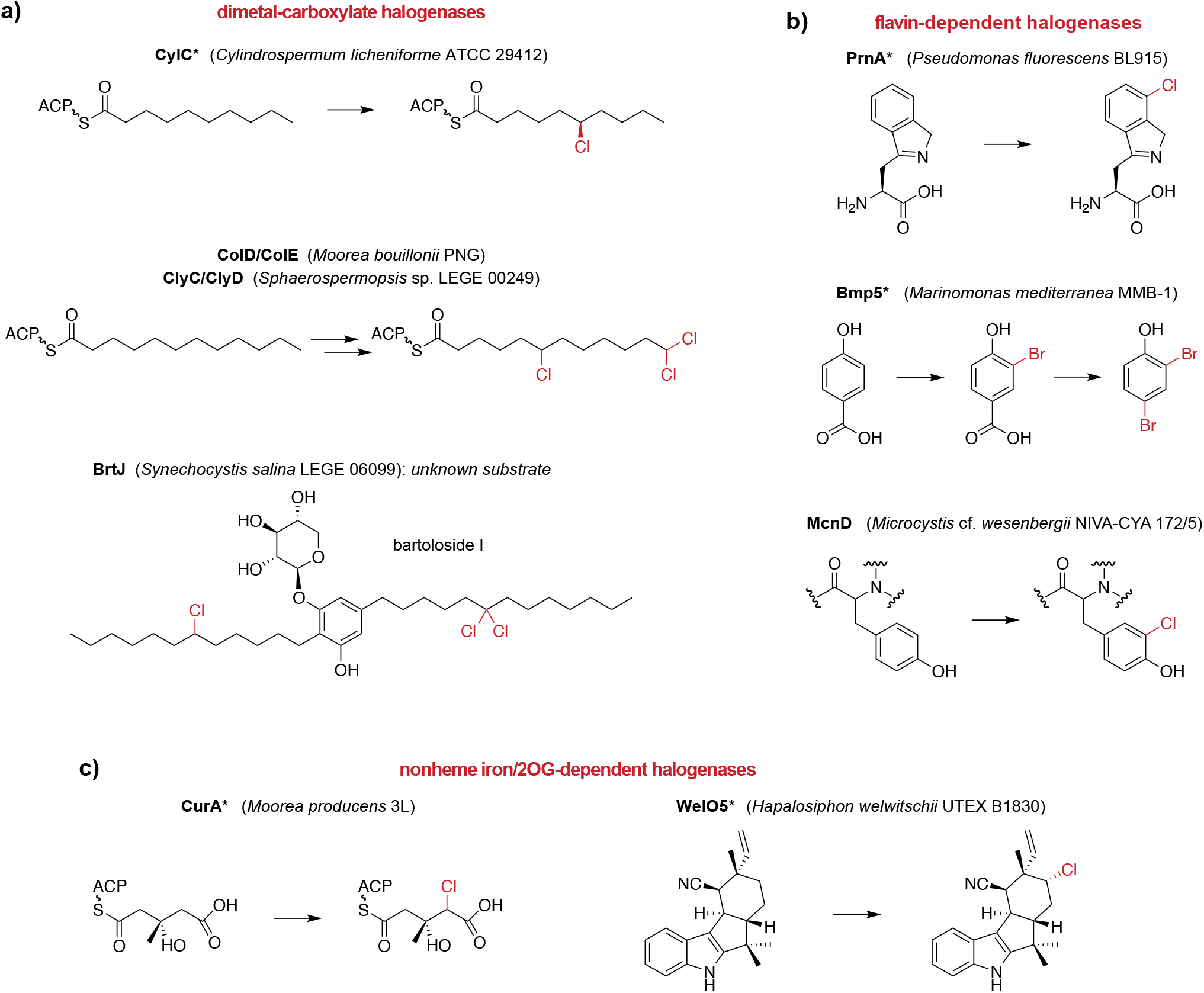
Selected examples of halogenation reactions catalyzed by different classes of microbial enzymes, with a focus on cyanobacterial halogenases. An asterisk denotes that the enzyme has been biochemically characterized. ACP – acyl carrier protein.

Recent efforts to characterize the biosynthesis of structurally unusual cyanobacterial natural products have uncovered a distinct class of halogenating enzymes. Using a genome mining approach, Nakamura et al. (2012) discovered the cylindrocyclophane biosynthetic gene cluster (BGC) in the cyanobacterium *Cylindrospermum licheniforme* ATCC 29412 [26]. The natural paracyclophane natural products were found to be assembled from two chlorinated alkylresorcinol units [27]. The paracyclophane macrocycle is created by forming two C-C bonds using a Friedel–Crafts-like alkylation reaction catalyzed by the enzyme CylK [27] (Fig. 1). Therefore, although many cylindrocyclophanes are not halogenated, their biosynthesis involves a halogenated intermediate [26, 27], a process termed a cryptic halogenation [28]. Nakamura et al. (2017) showed that the CylC enzyme was responsible for regio- and stereoselectively installing a chlorine atom onto the fatty acid-derived *sp*^*3*^ carbon center of a biosynthetic intermediate that is subsequently elaborated to the key alkylresorcinol monomer (Fig. 1). To date, CylC is the only characterized dimetal-carboxylate halogenase (this classification is based on both biochemical evidence and similarity to other diiron-carboxylate proteins) [27]. Homologs of CylC have been found in the BGCs of the columbamides [29], bartolosides [30], microginin [27], puwainaphycins/minutissamides [31], and chlorosphaerolactylates [32], all of which produce halogenated metabolites. CylC-type enzymes bear low sequence homology to dimetal desaturases and *N*-oxygenases [27], functionalize C-H bonds in aliphatic moieties at either terminal or mid-chain positions, and are likely able to carry out gem-dichlorination (Kleigrewe 2015, Leão 2015). The reactivity displayed by CylC and its homologs is of interest for biocatalysis, in particular because this type of carbon center activation is often inaccessible to organic synthesis [15, 33]. An understanding of the molecular basis for the halogenation of different positions and for chain-length preference will also be of value for biocatalytic applications. Hence, accessing novel variants of CylC enzymes will facilitate the functional characterization of this class of halogenases, mechanistic studies, and biocatalyst development.

Here, we provide an in-depth analysis of the diversity, distribution and context of CylC homologs in microbial genomes. Using both publicly available genomes and our in-house culture collection of cyanobacteria (LEGEcc), we report that CylC enzymes are common in cyanobacterial genomes, found in numbers comparable to those of flavin-dependent or nonheme iron/2OG-dependent halogenases. We additionally show that CylC homologs are distributed throughout the cyanobacterial phylogeny and are, to a great extent, part of cryptic BGCs with diverse architectures, underlining the potential for NP discovery associated with this new halogenase class.

## Methods

### Sequence similarity networks and Genomic Neighborhood Diagrams

Sequence similarity networks (SSNs) were generated using the EFI-EST sever, following a “Sequence BLAST” of CylC (AFV96137) as input [34], using negative log e-values of 2 and 40 for UniProt BLAST retrieval and SSN edge calculation, respectively. This SSN edge calculation cutoff was found to segregate the homologs into different SSN clusters, less stringent cutoff values resulted in a single SSN cluster. The 153 retrieved sequences and the query sequence were then used to generate the SSNs with an alignment score threshold of 42 and a minimum length of 90. The networks were visualized in Cytoscape (v3.80). The full SSN obtained in the previous step was used to generate Genomic Neighborhood Diagrams (GNDs) using the EFI-GNT tool [34]. A Neighborhood Size of 10 was used and the Lower Limit for Co-occurrence was 20%. The resulting GNDs were visualized in Cytoscape (Fig. 2).

**Figure 2.**
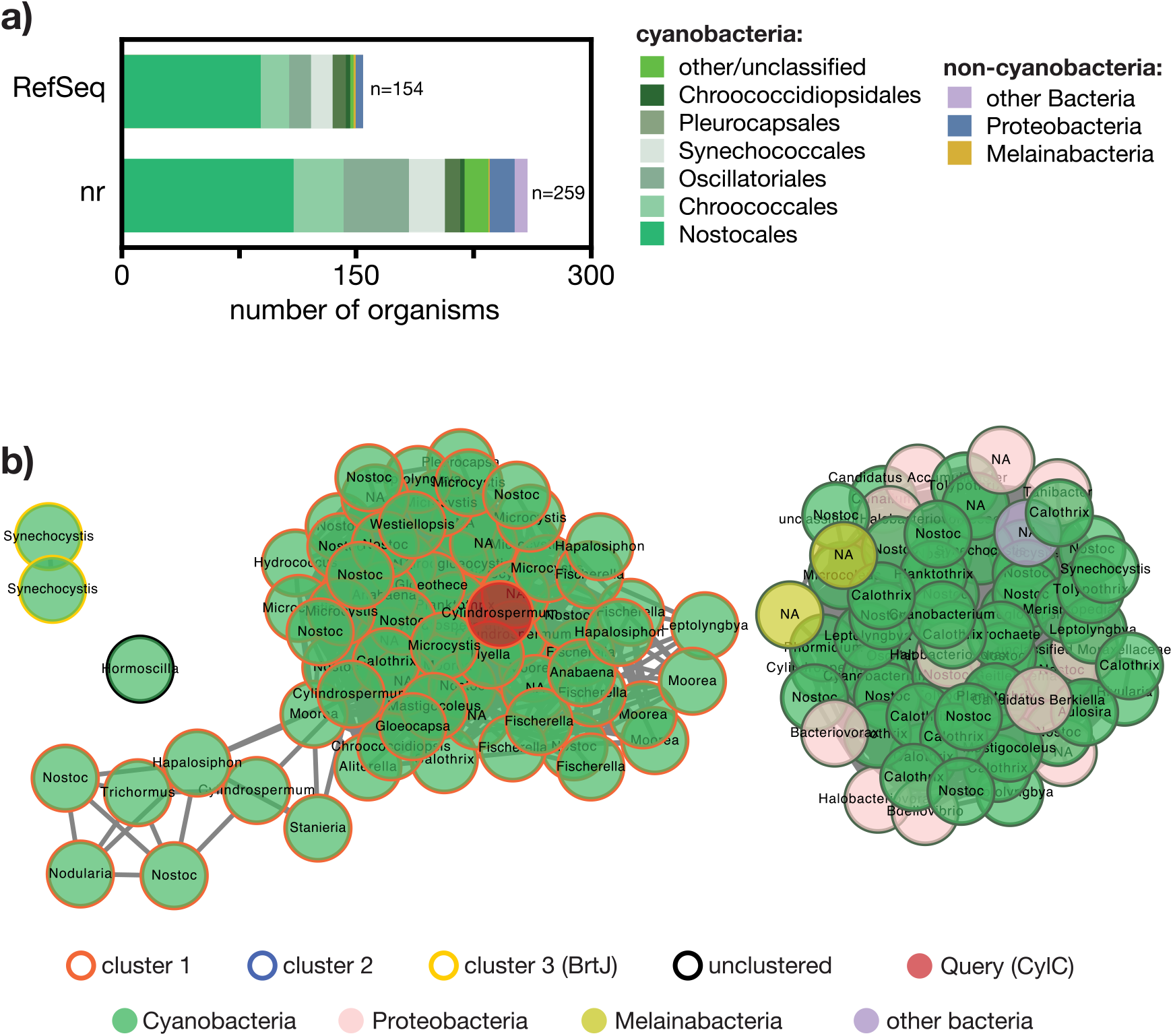
Abundance of CylC homologs in bacteria. a) BLASTp using CylC (GenBank accession no: ARU81117) as query against different databases, shows that these dimetal-carboxylate enzymes are found almost exclusively in cyanobacteria. b) Sequence Similarity Network (SSN) of CylC depicting the similarity-based clustering of UniProt-derived protein sequences with homology (BLAST e-value cutoff 1×10^−2^, edge e-value cutoff 1×10^−40^) to CylC (GenBank accession no: ARU81117). In each node, the bacterial genus for the corresponding UniProt entry is shown (NA – not attributed).

### Cyanobacterial strains and growth conditions

Freshwater and marine cyanobacteria strains from Blue Biotechnology and Ecotoxicology Culture Collection (LEGEcc) (CIIMAR, University of Porto) were grown in 50 mL Z8 medium [35] or 50 mL Z8 25‰ sea salts (Tropic Marine) with vitamin B12, with orbital shaking (∼200 rpm) under a regimen of 16 h light (25 μmol photons m-2 s -1)/8 h dark at 25 °C.

### Genomic DNA extraction

Fifty milliliters of each cyanobacterial strain were centrifuged at 7000 ×*g* for 10 min. The cell pellets were used for genomic DNA (gDNA) extraction using the PureLink ® Genomic DNA Mini Kit (Thermo Fisher Scientific®) or NZY Plant/Fungi gDNA Isolation kit (Nzytech), according to the manufacturer’s instructions.

### Primer design

Basic local alignment search tool (BLAST) searches using CylC [*Cylindrospermum licheniforme* UTEX B 2014] as query identified related genes (for tBLASTn: 31-93% amino acid identity). We discarded nucleotide hits with a length <210 and e-values <1×10^−10^. The complete sequences (56 *cylC* homolog sequences, Table S1) were collected from NCBI and aligned using MUltiple Sequence Comparison by Log-Expectation (MUSCLE) [36]. Phylogenetic analysis of the hits was performed using FastTree GTR with a rate of 100. *Streptomyces thioluteus aurF*, encoding a distant dimetal-carboxylate protein [27] was used as an outgroup (AJ575648.1:4858-5868). We divided the phylogeny of *cylC* homologs in five groups with moderate similarity (Fig. S1). The regions of higher similarity within each group were selected for degenerate primer design (Table 1).

**Table 1.**
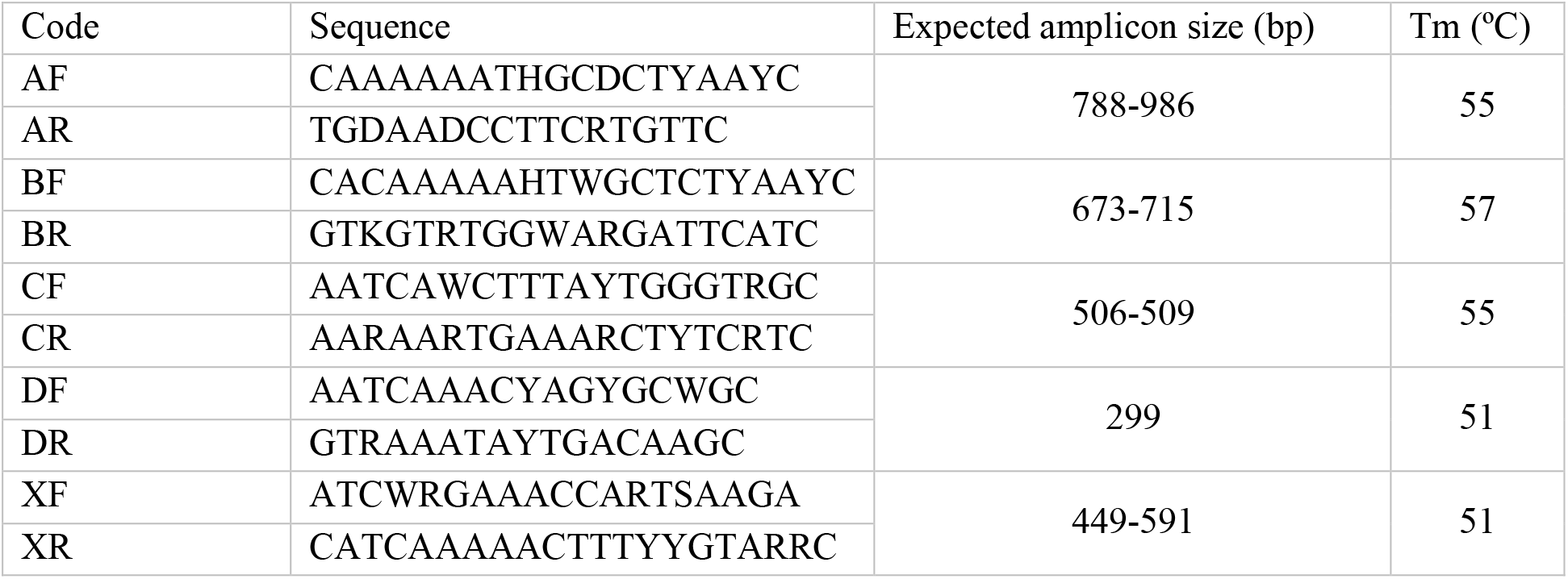
Degenerate primers

### PCR conditions

The PCR to detect *cylC* homologs were conducted in a final volume of 20 µL, containing 6.9 µL of ultrapure water, 4.0 µL of 5× GoTaq Buffer (Promega), 2.0 µL of MgCl_2_, 1.0 µL of dNTPs, 2.0 µL of reverse and 2.0 µL of forward primer (each at 10 µM), 0.1 µL of GoTaq and 2.0 µL of cyanobacterial gDNA. PCR thermocycling conditions were: denaturation for 5 min at 95 °C; 35 cycles with denaturation for 1 min at 95 °C, primer annealing for 30 s at different temperatures (55 °C for group A; 57°C for group B; 55 °C for group C; 51 °C for group D; 51 °C for group X) and extension for 1 min at 72 °C; and final extension for 10 min at 72 °C.

When not already available, the 16S rRNA gene for a tested strain was amplified by PCR, using standard primers for amplification (CYA106F 5’ CGG ACG GGT GAG TAA CGC GTG A 3’ and CYA785R 5’ GAC TAC WGG GGT ATC TAA TCC 3’). The PCR reactions were conducted in a final volume of 20 µL, containing 6.9 µL of ultrapure water, 4.0 µL of 5× GoTaq Buffer, 2.0 µL of MgCl_2_, 1.0 µL of dNTPs, 2.0 µL of primer reverse and 2.0 µL of primer forward (each one at 10 µM), 0.1 µL of GoTaq and 2.0 µL of cyanobacterial DNA. PCR thermocycling conditions were: denaturation for 5 min at 95 °C; 35 cycles with denaturation for 1 min at 95 °C, primer annealing for 30 s at 52 °C and extension for 1 min at 72 °C; and final extension for 10 min at 72 °C. Amplicon sizes were confirmed after separation in a 1.0% agarose gel.

### Cloning and sequencing

The *cylC* homolog and 16S rRNA gene sequences were obtained either directly from the NCBI or through sequencing. To obtain high quality sequences, the TOPO PCR cloning (Invitrogen) was used. The TOPO cloning reaction was conducted in a final volume of 3 µL, containing 1 µL of fresh PCR product, 1 µL of salt solution, 0.5 µL of TOPO vector and 0.5 µL of water. The reaction was incubated for 20 min at room temperature. Three-microliters of TOPO reaction were added into a tube containing chemically competent *E. coli* (Top10, Life Technologies) cells. After 30 min of incubation on ice, the cells were placed for 30 s at 42 °C without shaking and were then immediately transferred to ice. 250 µL of room temperature SOC medium were added to the previous mixture and the tube was horizontally shaken at 37 °C for 1 h (180rpm). 60 µL of the different cloning reactions were spread onto LB ampicillin/X-gal plates and incubated overnight at 37 °C.

Two or three positive colonies from each reaction were tested by colony-PCR. The PCR was conducted in a final volume of 20 µL, containing 10.9 µL of ultrapure water, 4.0 µL of 5x GoTaq Buffer, 2.0 µL of MgCl_2_, 1.0 µL of dNTPs, 1.0 µL of reverse pUCR and 1.0 µL of forward pUCF primers (each at 20 µM), 0.1 µL of GoTaq and the target colony. PCR thermocycling conditions were: denaturation for 5 min at 95 °C; 35 cycles with denaturation for 1 min at 95 °C, primer annealing for 30 s at 50 °C and extension for 1 min at 72 °C; and final extension for 10 min at 72 °C. Amplicon sizes were confirmed after separation in an 1.0 % agarose gel. Selected colonies were incubated overnight at 37 °C (180 rpm), in 5 mL of LB supplemented with 100 µg mL^-1^ ampicillin. The plasmids containing the amplified PCR products were extracted (NZYMiniprep kits) and Sanger sequenced using pUC primers.

### Cyanobacteria genome sequencing

Many of the LEGEcc strains are non-axenic, and so before extraction of gDNA for genome sequencing, an evaluation of the amount of heterotrophic contaminant bacteria in cyanobacterial cultures was performed by plating onto Z8 or Z8 with added 2.5% sea salts (Tropic Marine) and vitamin B_12_ (10 µg/L) agar medium (depending the original environment) supplemented with casamino acids (0.02% wt/vol) and glucose (0.2% wt/vol) [37]. The plates were incubated for 2-4 days at 25 °C in the dark and examined for bacterial growth. Those cultures with minimal contamination were used for DNA extraction for genome sequencing. The selection of DNA extraction methodology used was based on morphological features of each strain. Total genomic DNA was isolated from a fresh or frozen pellet of 50 mL culture using a CTAB-chloroform/isoamyl alcohol-based protocol [38] or using the commercial PureLink Genomic DNA Mini Kit (Thermo Fisher Scientific®) or the NZY Plant/Fungi gDNA Isolation kit (NZYTech). The latter included a homogenization step (grinding cells using a mortar and pestle with liquid nitrogen) before extraction using the standard kit protocol. The quality of the gDNA was evaluated in a DS-11 FX Spectrophotometer (DeNovix) and 1 % agarose gel electrophoresis, before genome sequencing, which was performed elsewhere (Era7, Spain and MicrobesNG, UK) using 2 × 250 bp paired-end libraries and the Illumina platform (except for *Synechocystis* sp. LEGE 06099, whose genome was sequenced using the Ion Torrent PGM platform). A standard pipeline including the identification of the closest reference genomes for reading mapping using Kraken 2 [39] and BWA-MEM to check the quality of the reads [40] was carried out, while *de novo* assembly was performed using SPAdes [41]. The genomic data obtained for each strain was treated as a metagenome. The contigs obtained as previously mentioned were analyzed using the binning tool MaxBin 2.0 [42] and checked manually in order to obtain only cyanobacterial contigs. The draft genomes were annotated using the NCBI Prokaryotic Genome Annotation Pipeline (PGAP) [43] and submitted to GenBank under the BioProject number SUB8150995. In the case of *Hyella patelloides* LEGE 07179 and *Sphaerospermopsis* sp. LEGE 00249 the assemblies had been previously deposited in NCBI under the BioSample numbers SAMEA4964519 and SAMN15758549, respectively.

### Genomic context of CylC homologs

BLASTp searches using CylC [*Cylindrospermum licheniforme* UTEX B 2014] as query identified related CylC homologs within the publicly available cyanobacterial genomes and in the genomes of LEGEcc strains. We annotated the genomic context for each CylC homolog using antiSMASH v5.0 [44] and manual annotation through BLASTp of selected proteins. Some BGCs were not identified by antiSMASH and were manually annotated using BLASTp searches.

### Phylogenetic analysis

Nucleotide sequences of *cylC* homologs obtained from the NCBI and from genome sequencing in this study, were aligned using MUSCLE from within the Geneious R11.0 software package (Biomatters). The nucleotide sequence of the distantly-related dimetal-carboxylate protein AurF [27] from *Streptomyces thioluteus* (AJ575648.1:4858-5868) was used as an outgroup. The alignments, trimmed to their core 788, 673, 506, 299 and 499 positions (for group A, B, C, D and X, respectively), were used for phylogenetic analysis, which was performed using FastTree 2 (from within Geneious), using a GTR substitution model (from jmodeltest, [45]) with a rate of 100 (Fig. S2).

For the phylogenetic analysis based on the 16S rRNA gene (Fig. 3, Fig. S3), the corresponding nucleotide sequences were retrieved from the NCBI (from public available genomes until March 16, 2020) or from sequence data (amplicon or genome) obtained in this study. The sequences were aligned as detailed for *cylC* homologs and trimmed to the core shared positions (663). A RAxML-HPC2 phylogenetic tree inference using maximum likelihood/rapid bootstrapping run on XSEDE (8.2.12) with 1000 bootstrap iterations in the Cipres platform [46] was performed.

**Figure 3.**
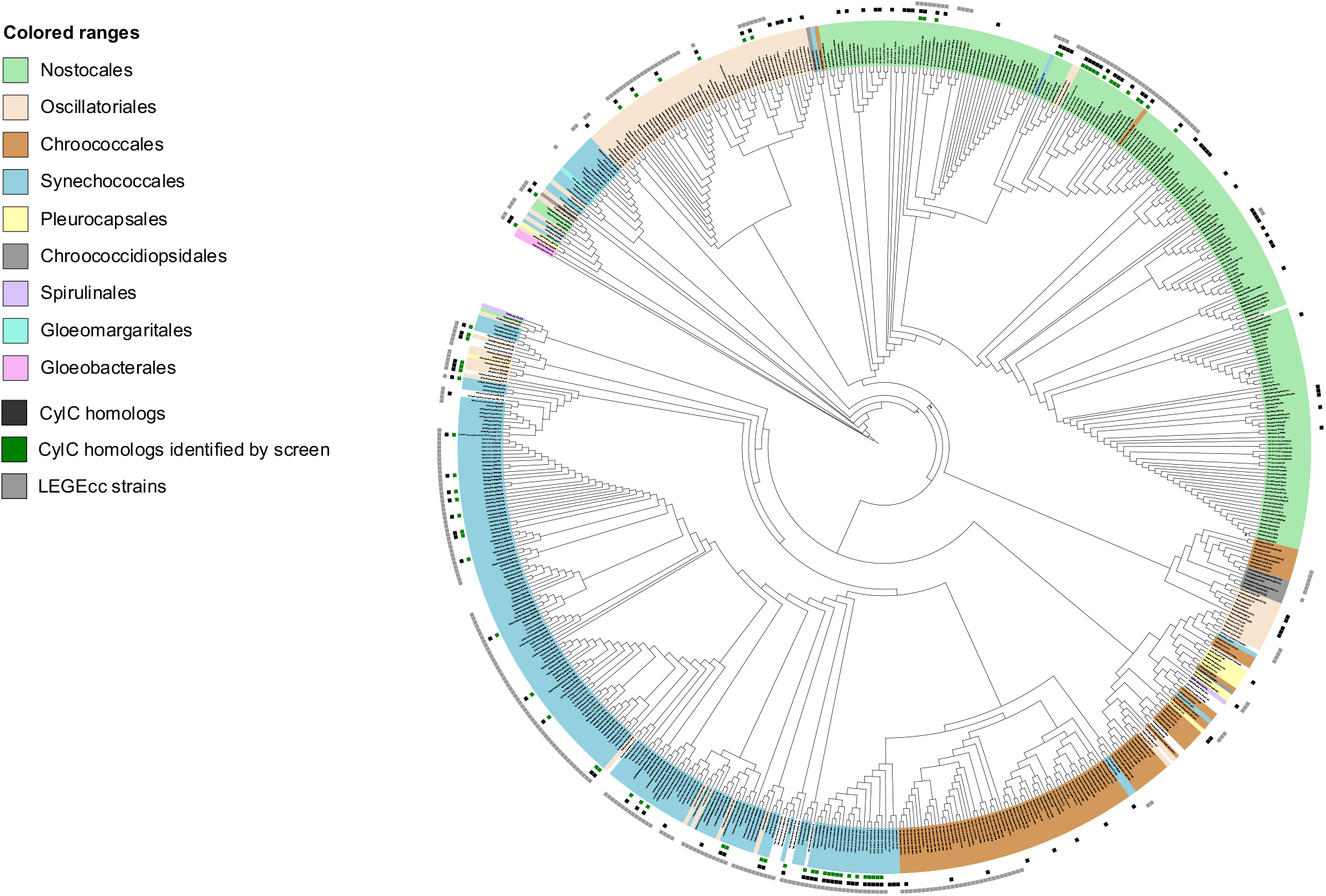
RAxML cladogram of the 16S rRNA gene of LEGEcc strains (grey squares) and from cyanobacterial strains with NCBI-deposited reference genomes, screened in this study. Taxonomy is presented at the order level (colored rectangles). Strains whose genomes encode CylC homologs are denoted by black squares. Green squares indicate that at least one homolog was detected by PCR-screening and verified by retrieving the sequence of the corresponding amplicon by cloning followed by Sanger sequencing. *Gloeobacter violaceus* PCC 7421 served as an outgroup. A version of this cladogram including the bootstrap values for 1000 replications is provided as Supplementary Material.

The amino acid sequences of CylC homologs were aligned using MUSCLE from within the Geneious software package (Biomatters). The alignments were trimmed to their core 333 residues and used for phylogenetic analysis, which was performed using RAxML-HPC2 phylogenetic tree inference using maximum likelihood/rapid bootstrapping run on XSEDE (8.2.12) with 1000 bootstrap iterations in the Cipres platform [46] (Fig. 4c).

**Figure 4.**
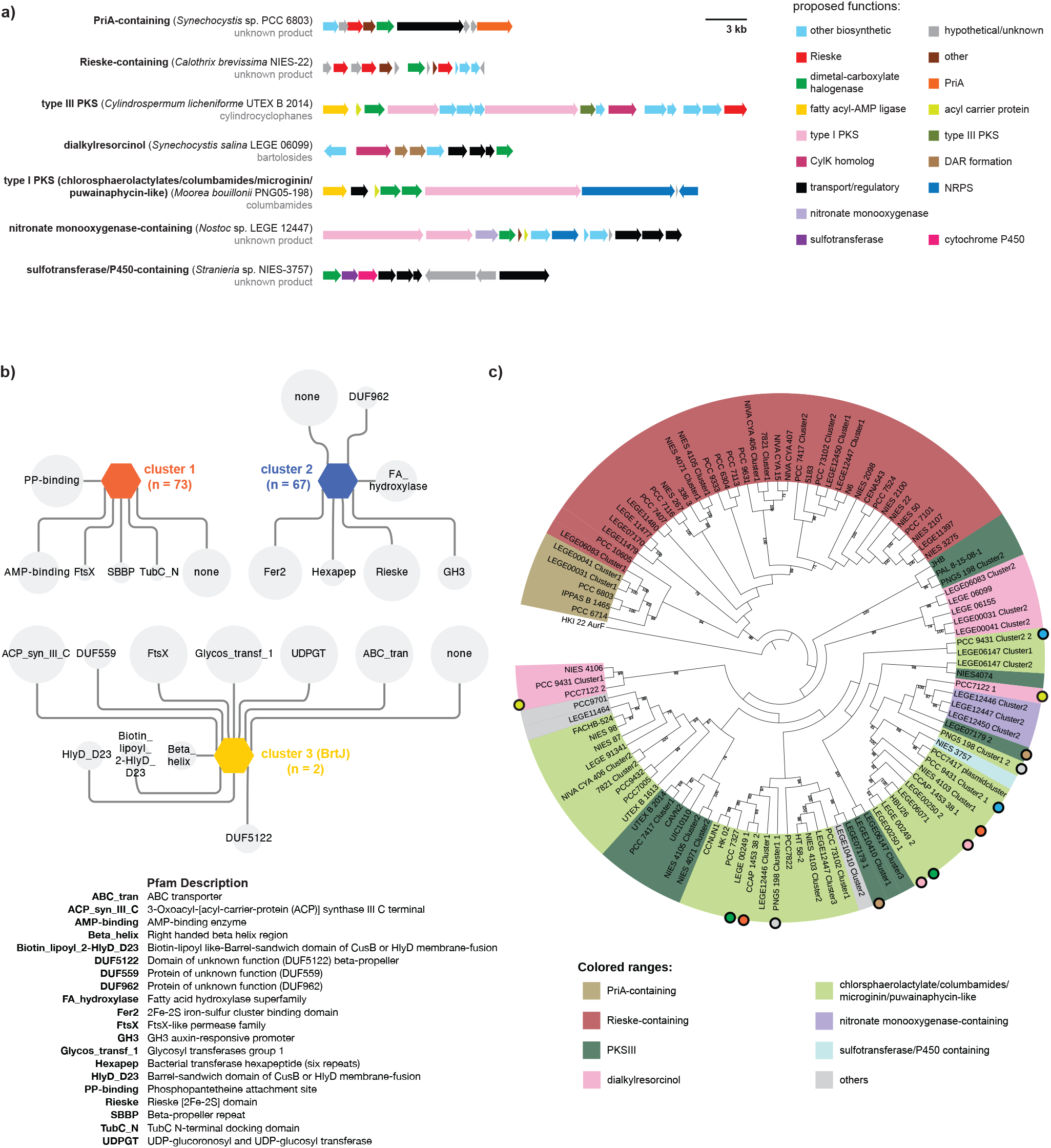
Diversity and genomic context of CylC-like enzymes BGCs. a) Examples of the different BGCs architectures found among the clusters encoding CylC homologs. b) Genome Neighborhood Diagram (GND) depicting the Pfam domains associated with each cluster from the initial SSN of CylC homologs. The size of each node is proportional to the prevalence of the Pfam domain within the genomic context of the CylC homologs from each SSN cluster. c) RAxML cladogram (1000 replicates, shown are bootstrap values > 70%) of CylC homologs. The different colors represent a categorization based on common genes found within the associated biosynthetic gene clusters (see legend). Circles of the same color depict CylC homologs encoded by the same BGC. AurF (*Streptomyces thioluteus* HKI-22) was used as an outgroup.

### CORASON analysis

CORASON, a bioinformatic tool that computes multi-locus phylogenies of BGCs within and across gene cluster families [47], was used to analyze cyanobacterial genomes collected from the NCBI and the LEGEcc genomes (Table S2). In total 2059 cyanobacterial genomes recovered from NCBI and 56 additional LEGE genomes were used in the analysis. The amino acid sequences of CurA (AAT70096.1), WelO5 (AHI58816.1), McnD (CCI20780.1), Bmp5 (WP_008184789.1), PrnA (WP_044451271.1) and CylC (ARU81117.1) were used as query and, for each enzyme, a reference genome was selected (Table S2). To increase the phylogenetic resolution, selected genomes were removed from the analysis of enzymes CylC, PrnA, CurA, McnD and Bmp5 (Table S2). Additionally, for the CylC analysis, a few BGCs were manually extracted and included in the analysis (Table S2) since they were not detected by CORASON.

### Prevalence of halogenases in cyanobacterial genomes

Representative proteins of each class were used as query in each search: CylC (ARU81117.1), BrtJ (AKV71855.1), “Mic” (WP_002752271.1) - the halogenase in the putative microginin gene cluster – ColD (AKQ09581.1), ColE (AKQ09582.1), NocO (AKL71648.1), NocN (AKL71647.1) for dimetal-carboxylate halogenases; PrnA (WP_044451271.1), Bmp5 (WP_008184789.1), and McnD (CCI20780.1) for flavin-dependent halogenases; the halogenase domains from CurA (AAT70096.1), and the halogenases Barb1 (AAN32975.1), HctB (AAY42394.1), WelO5 (AHI58816.1) and AmbO5 (AKP23998.1) for nonheme iron-dependent halogenases). Non-redundant sequences obtained for these searches using a 1×10^−20^ e-value cutoff, which represents a percentage identity between the query and target protein superior to 30%, were considered to share the same function as the query.

## Results and Discussion

### CylC-like halogenases are mostly found in cyanobacteria

To investigate the distribution of CylC homologs encoded in microbial genomes, we first searched the reference protein (RefSeq) or non-redundant protein sequences (nr) databases (NCBI) for homologs of CylC or BrtJ, using the Basic Local Alignment Search Tool, BLASTp (min 25% identity, 9.9×10^−20^ E-value and 50% coverage). A total of 128 and 246 homologous unique protein sequences were retrieved using the RefSeq or nr databases, respectively; in both cases, sequences were primarily from cyanobacteria (96 and 88%, respectively) (Fig. 2a). We then used the Enzyme Similarity Tool of the Enzyme Function Initiative (EFI-EST) [34] to evaluate the sequence landscape of dimetal-carboxylate halogenases. Using CylC as query, we obtained a SSN (sequence similarity network) composed of 154 sequences retrieved from the UniProt database [48] (Fig. 2b). The SSN featured two major clusters, one containing homologs from diverse cyanobacterial genera, the other composed of homologs from several cyanobacteria, with a few from proteobacteria (mostly deltaproteobacteria) and two from the cyanobacteria sister-phylum Melainabacteria. A third SSN cluster was composed only by the previously reported BrtJ enzymes and, finally, a homolog from the cyanobacterial genus *Hormoscilla* remained unclustered. We were unable to recover any SSN that included clusters containing other characterized enzyme functions, which attests to the uniqueness of the dimetal-carboxylate halogenases in the current protein-sequence landscape.

### CylC homologs are widely distributed throughout the phylum Cyanobacteria

With the intent of accessing a wide diversity of CylC homolog sequences, we decided to use a degenerate-primer PCR strategy to discover additional homologs in cyanobacteria from the LEGEcc culture collection [49], because the phylum Cyanobacteria is diverse and still underrepresented in terms of genome data [50-55]. The LEGEcc culture collection maintains cultures isolated from diverse freshwater and marine environments, mostly in Portugal, and, for example, contains all known bartoloside-producing strains [30]. Primers were designed based on 54 nucleotide sequences retrieved from the NCBI that were selected to represent the phylogenetic diversity of CylC homologs (Fig. S1). Due to the lack of highly conserved nucleotide sequences among all homologs considered, we divided the nucleotide alignment into five groups and designed a degenerate primer pair for each. Upon screening 326 strains from LEGEcc using the five primer pairs, we retrieved 89 sequences encoding CylC homologs, confirmed through cloning and Sanger sequencing of the obtained amplicons. We were unable to directly analyze the diversity of the entire set of LEGEcc-derived *cylC* amplicons due to low overlap between sequences obtained with different primers. As such, we performed a phylogenetic analysis of the diversity retrieved with each primer pair (Fig. S2), by aligning the PCR-derived sequences with a set of diverse *cylC* genes retrieved from the NCBI. For some strains, our PCR screen retrieved more than one homolog using different primer pairs (e.g. *Nostoc* sp. LEGE 12451 or *Planktothrix mougeotii* LEGE 07231). In general, and for each primer pair, the PCR screen retrieved mostly sequences that were closely related and associated to one or two phylogenetic clades. This can likely be explained by the geographical bias that might exist in the LEGEcc culture collection [49] and/or with primer design and PCR efficiency issues, which might have favored certain phylogenetic clades.

To access full-length sequences of the CylC homologs identified among LEGEcc strains, as well as their genomic context, we undertook a genome-sequencing effort informed by our PCR screen. We selected 21 strains for genome sequencing, which represents the diversity of CylC homologs observed in the different PCR screening groups. The resulting genome data was used to generate a local BLAST database and the homologs were located within the genomes. In some cases, additional homologs that were not detected in the PCR screen were identified. Overall, 33 full-length genes encoding CylC homologs were retrieved from LEGEcc strains.

To explore the phylogenetic distribution of CylC homologs encoded in publicly available reference genomes and the herein sequenced LEGEcc genomes, we aligned the 16S rRNA genes from 648 strains with RefSeq genomes and the LEGEcc strains that were screened by PCR in this study. Using this dataset, we performed a phylogenetic analysis which indicated that CylC homologs are broadly distributed through five Cyanobacterial orders: Nostocales, Oscillatoriales, Chroococcales, Synechococcales and Pleurocapsales (Fig. 3, Fig. S3). It is noteworthy that the cyanobacterial orders for which we did not find CylC homologs (Chroococcidiopsidales, Spirulinales, Gloeomargaritales and Gloeobacterales) are poorly represented in our dataset (Fig. 3, Fig. S3). However, our previous BLASTp search against the nr database did retrieve two close homologs in two Chroococcidiopsidales strains (genera *Aliterella* and *Chroococcidiopsis*) and a more distant homolog in a *Gloeobacter* strain (Gloeobacterales) (Table S3). Given the wide but punctuated presence of CylC homologs among the cyanobacterial diversity considered in this study, it is unclear how much of the current CylC homolog distribution reflects vertical inheritance or horizontal gene transfer events.

### Diversity of BGCs encoding CylC homologs

To characterize the biosynthetic diversity of BGCs encoding CylC homologs, which were found in 78 cyanobacterial genomes (21 from LEGEcc and 57 from RefSeq) from different orders, we first submitted these genome sequences for antiSMASH [44] analysis. 55 CylC-encoding BGCs were detected, which were classified as resorcinol, NRPS, PKS, or hybrid NRPS-PKS. Given the number of CylC homolog-encoding genes detected in these genomes (105), we considered that several BGCs might have not been identified with antiSMASH. Therefore, we performed manual annotation of the genomic contexts of the CylC homologs and were able to identify 20 additional BGCs. Upon analysis of the entire set of CylC-encoding BGCs, we classified the BGCs in seven major categories, based on their overall architecture, which we designated as follows (listed in decreasing abundance): Rieske-containing (n = 36), type I PKS (chlorosphaerolactylate/columbamide/microginin/puwainaphycin-like, n = 29), type III PKS (n = 13), dialkylresorcinol (n = 8), PriA-containing (n = 5), nitronate monooxygenase-containing (n = 3) and cytochrome P450/sulfotransferase-containing (n = 1) (Fig. 4a, Figs. S4-S10). Three BGCs were excluded from our classification since they were only partially sequenced (Fig. S11). Examples of each of the cluster architectures are presented in Fig. 4a and schematic representations of each of the 98 classified BGCs are presented in Supplementary Figures S4-S10. It should be stressed that within several of these seven major categories, there is still considerable BGC architecture diversity, notably within the dialkylresorcinol, type I and type III PKS BGCs. Rieske-containing BGCs are not associated with any known NP and encode between two and four proteins with Rieske domains. Most contain a sterol desaturase family protein, feature a single CylC homolog and are chiefly found among Nostocales and Oscillatoriales (Fig. S4). PriA-containing BGCs encode, apart from the Primosomal protein N’ (PriA), a set of additional diguanylate cyclase/phosphodiesterase, aromatic ring-hydroxylating dioxygenase subunit alpha and a ferritin-like protein and were only detected in *Synechocystis* spp. (Fig. S5). These are similar to the Rieske-containing BGCs; however, in strains harboring PriA-containing BGCs, the additional functionalities that are found in the Rieske-containing BGCs can be found dispersed throughout the genome (Table S4). In our dataset, a single sulfotransferase/P450 containing BGC was detected in *Stanieria* sp. and was unrelated to the above-mentioned architectures (Fig. S6). Type I PKS BGCs encode clusters similar to those of the chlorosphaerolactylates, columbamides, microginins and puwainaphycins and typically feature a fatty acyl-AMP ligase (FAAL) and an acyl carrier protein upstream of one or two CylC homologs and a type I PKS downstream of the CylC homolog(s). These were found in Nostocales and Oscillatoriales strains (Fig. S7). Taken together with the known NP structures associated with these BGCs [29, 56, 57], we can expect that the encoded metabolites feature halogenated fatty acids in terminal or mid-chain positions. BGCs of the dialkylresorcinol type, which contain DarA and DarB homologs (Bode 2013, Leão 2015), including several bartoloside-like clusters (found only in LEGEcc strains), were detected in Nostocales, Pleurocapsales and Chroococcales (Fig. S8). Type III PKS BGCs encoding CylC homologs, which include a variety of cyclophane BGCs, were detected in the Nostocales, Oscillatoriales and Pleurocapsales (Fig. S9). Finally, nitronate monooxygenase-containing BGCs, which are not associated with any known NP, were only found in Nostocales strains from the LEGEcc and featured also genes encoding PKSI, ferredoxin, ACP or glycosyl transferase (Fig. S10).

A less BGC-centric perspective of the genomic context of CylC homologs could be obtained through the Genome Neighborhood Tool of the EFI (EFI-GNT, [58]). Using the previously generated SSN as input, we analyzed the resulting Genomic Neighborhood Diagrams (Fig. 4b), which indicated that the three SSN clusters had entirely different genomic contexts (herein defined as 10 upstream and 10 downstream genes from the *cylC* homolog). The SSN cluster that encompasses CylC and its closest homologs indicates that these enzymes associate most often with PP-binding (ACP/PCPs) and AMP-binding (such as FAALs) proteins. Regarding the SSN cluster that includes both cyanobacterial and non-cyanobacterial CylC homologs, their genomic contexts most prominently feature Rieske/[2Fe-2S] cluster proteins as well as fatty acid hydroxylase family enzymes. The cyanobacterial homologs are exclusively encoded in the Rieske and PriA-containing BGCs. Homologs from this particular SSN cluster may not require a phosphopantetheine tethered substratei+ as no substrate activation or carrier proteins/domains were found in their genomic neighborhoods, or may act on central fatty acid metabolism intermediates. The BrtJ SSN cluster, composed only of the two reported BrtJ enzymes, shows entirely different surrounding genes, obviously corresponding to the *brt* genes. Also noteworthy is the considerable number of proteins with unknown function found in the vicinity of dimetal-carboxylate halogenases, suggesting that uncharted biochemistry is associated with these enzymes.

Since SSN analysis generated only three clusters of CylC homologs, we next investigated the genetic relatedness among these enzymes and how it correlates to BGC architecture. We performed a phylogenetic analysis of the CylC homologs from the 98 classified and 3 unclassified BGCs (Fig. 4c). Our analysis indicated that PriA-containing and Rieske-containing BGCs formed a well-supported clade. Its sister clade contained homologs from the remaining BGCs. Within this larger clade, homologs associated with the type I PKS, dialkylresorcinol or type III PKS BGCs were found to be polyphyletic. In some cases, the same BGC contained distantly related CylC homologs (e.g. *Hyella patelloides* LEGE 07179, *Anabaena cylindrica* PCC 7122) (Figure 4c). This analysis also revealed that several strains (Fig. 5c) encode two or three phylogenetically distant CylC homologs in different BGCs. Overall, our data shows that CylC homologs have evolved to interact with different partner enzymes to generate chemical diversity, but that their phylogeny is, in some cases, not entirely consistent with BGC architecture. These observations suggest that functionally convergent associations between CylC homologs and other proteins have emerged multiple times during evolution. Examples include the CylC/CylK and BrtJ/BrtB associations, which use cryptic halogenation to achieve C-C and C-O bond formation, respectively [27, 59]. However, the role of the CylC homolog-mediated halogenation of fatty acyl moieties observed for other cyanobacterial metabolites is not currently understood. Interestingly, while a number of CylC homologs, including those that are part of characterized BGCs, likely act on ACP-tethered fatty acyl substrates [27, 59], those from the PriA-Rieske- and cytochrome P450/sulfotransferase categories do not have a neighboring carrier protein and therefore might not require a tethered substrate. This would be an important property for a CylC-like biocatalyst [15].

**Figure 5.**
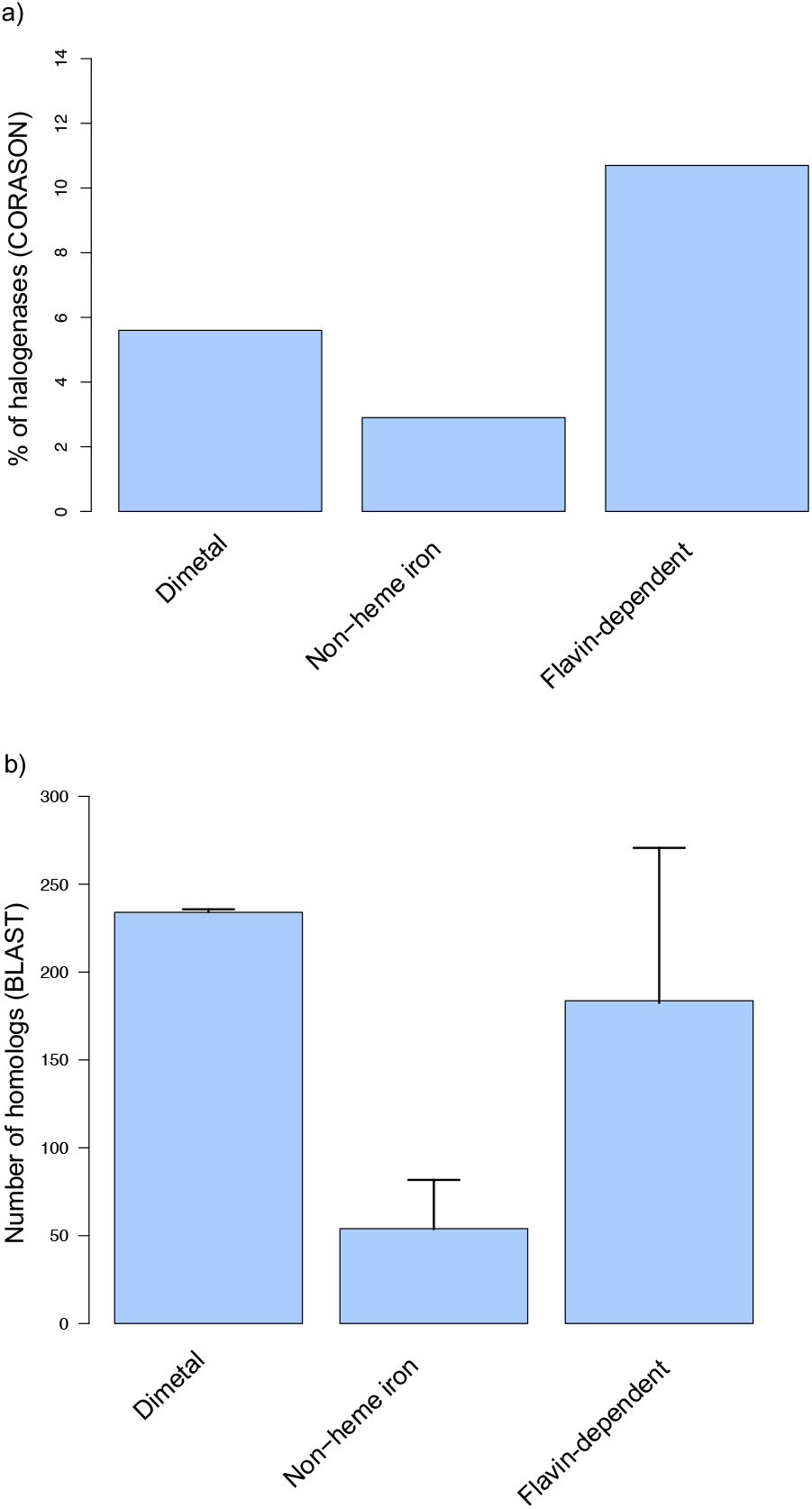
Prevalence of cyanobacterial halogenases. Frequency of halogenases in Cyanobacteria from CORASON analysis (A) and NCBI BLASTp analysis (B). (A) Dimetal-carboxylate halogenases: CylC - NCBI reference genomes, n = 2054 and LEGEcc genomes, n = 41 CylC-containing BGCs and 56 genomes; Flavin-dependent halogenases: PrnA - NCBI reference genomes, n = 2051 and LEGEcc genomes, n = 56 genomes; Bmp5-NCBI reference genomes, n = 2050 and LEGEcc genomes, n = 56 genomes; McnD: NCBI reference genomes, n = 2052 and LEGEcc genomes, n = 54 genomes); Nonheme iron/2OG-dependent halogenases: halogenase domain from CurA - NCBI reference genomes, n = 2052 and LEGEcc genomes, n = 56 genomes. (B) Average of the total number of homologs per dimetal-carboxylate halogenases (CylC, BrtJ, “Mic”, ColD, ColE, NocO, NocN), flavin-dependent halogenases (Tryptophan 7-halogenase PrnA, Bmp5 and McnD) and nonheme iron/2OG-dependent halogenases (Barb1, HctB, WelO5, AmbO5 and the halogenase domain from CurA).

### CylC enzymes and other cyanobacterial halogenases

We sought to understand how CylC-type halogenases compare to other halogenating enzyme classes found in cyanobacteria in terms of prevalence and association with BGCs. To this end, we carried out a CORASON [47] analysis of publicly available cyanobacterial genomes (including non-reference genomes) and the herein acquired genome data from LEGEcc strains (a total of 2,115 cyanobacterial genomes). We used different cyanobacterial halogenases as input, namely CylC, McnD, PrnA, Bmp5, the 2OG-Fe(II) oxygenase domains from CurA and BarB1. CORASON attempts to retrieve genome context by exploring gene cluster diversity linked to enzyme phylogenies [47]. The CORASON analysis retrieved 117 (5.6%) dimetal-carboxylate halogenases, 61 (2.9%) nonheme iron-dependent halogenases and 226 (10.7%) flavin dependent halogenases from the cyanobacterial genomes (Fig. 5a). Using the protein homologs detected in BGCs by CORASON, a sequence alignment was performed for dimetal-carboxylate, nonheme iron/2OG-dependent and flavin-dependent halogenases. For nonheme iron/2OG-dependent halogenases, we excised the halogenase domain from multi-domain enzyme sequences. After removing repeated sequences and trimming the alignments to their core shared positions, maximum-likelihood phylogenetic trees were constructed for each halogenase class and BGCs were annotated manually (Figs. S12-S14). Flavin-dependent halogenases were commonly associated with cyanopeptolin, 2,4-dibromophenol and pyrrolnitrin BGCs and with orphan BGCs of distinct architectures (Fig. S12). Regarding nonheme iron/2OG-dependent halogenases, we identified barbamide, curacin, hectochlorin and terpene/indole [60] BGCs and several distinct orphan BGCs (Fig. S13). For dimetal-carboxylate halogenases, columbamide, microginin, chlorosphaerolactylate, bartoloside and cyclophane BGCs were identified (Fig. S14). However, while some of the CylC homolog-encoding orphan BGCs previously identified by antiSMASH and manual searches were detected by CORASON, the Rieske- and the PriA-containing BGCs were not. Hence, several CylC homologs were not accounted for in this analysis. For the same reasons, the other two halogenase types could also be missing some of its members in the CORASON-derived datasets. To circumvent this limitation and obtain a more comprehensive picture of the abundance of the three types of halogenase in cyanobacterial genomes, we used BLASTp searches against available cyanobacterial genomes in the NCBI database (including non-reference genomes). Several representatives of each halogenase class were used as query in each search (CylC, BrtJ, “Mic” – the halogenase in the putative microginin gene cluster – ColD, ColE, NocO and NocN for dimetal-carboxylate halogenases; PrnA, Bmp5 and McnD for flavin dependent halogenases; the halogenase domain from CurA and the halogenases BarB1, HctB, WelO5 and AmbO5 for nonheme iron-dependent halogenases). Non-redundant sequences obtained for these searches using a 1×10^−20^ e-value cutoff (corresponding to >30% sequence identity) were considered to share the same function as the query. It is worth mentioning that, for nonheme iron/2OG-dependent enzymes, a single amino acid difference can convert hydroxylation activity into halogenation [61], so it is possible that – at least for this class – the sequence space considered does not correspond exclusively to halogenation activity. Dimetal-carboxylate and flavin-dependent halogenase homologs were found to be the most abundant in cyanobacteria, each with roughly 0.2 homologs per genome, while nonheme iron/2OG-dependent halogenase homologs are less common (∼0.05 per genome) (Fig. 5b). Overall, our analyses indicate that homologs of each of the three halogenase classes are associated with a large number of orphan BGCs and represent opportunities for NP discovery. Particularly noteworthy, CylC-like enzymes are clearly a major group of halogenases in cyanobacteria, despite having been the latest to be discovered [27].

## Conclusion

The discovery of a new biosynthetic enzyme class brings with it tremendous possibilities for biochemistry and catalysis research, both fundamental and applied. Their functional characterization can also be used as a handle to identify and deorphanize BGCs that encode their homologs. CylC typifies an unprecedented halogenase class, which is almost exclusively found in cyanobacteria. By searching CylC homologs in both public databases and our in-house culture collection, we report here more than 100 new cyanobacterial CylC homologs. We found that dimetal-carboxylate halogenases are widely distributed throughout the phylum. The genomic neighborhoods of these halogenases are diverse and we identify a number of different BGC architectures associated with either one or two CylC homologs that can serve as starting points for the discovery of new NP scaffolds. In addition, the herein reported diversity and biosynthetic contexts of these enzymes will serve as a roadmap to further explore their biocatalysis-relevant activities. Finally, bartoloside-like BGCs and a CylC-associated BGC architecture (nitronate monooxygenase-containing) were found only in the LEGEcc, reinforcing the importance of geographically focused strain isolation and maintenance efforts for the Cyanobacteria phylum.

## Acknowledgments

We thank Hitomi Nakamura, Samantha Cassell, Diana Sousa and João Reis for technical assistance during this study, and the Blue Biotechnology and Ecotoxicology Culture Collection (LEGEcc) for the genomic DNA used for the PCR screening.

