## Supplementary Material for "Distribution and diversity of dimetal-carboxylate halogenases in cyanobacteria"

**Table S1.** Accession number of *cylC* homologs and *aurF* genes used for primer design.

| Name | Acession number |
|---|---|
| *Anabaena cylindrica* PCC 7122 | AP018166.1:4354075-4355373 |
| *Anabaena cylindrica* PCC 7122 | CP003659.1:3493241-3494452 |
| *Anabaena minutissima* UTEX B 1613 | MH325199.1:28248-29645 |
| *Aulosira laxa* NIES-50 | AP018307.1:7246161-7247336 |
| *Calothrix brevissima* NIES-22 | AP018207.1:8432194-8433369 |
| *Calothrix parasitica* NIES-267 | AP018227.1:c657283-656117 |
| *Calothrix* sp. 336/3 | CP011382.1:4155917-4157092 |
| *Calothrix* sp. NIES-2098 | AP018172.1:4832715-4833896 |
| *Calothrix* sp. NIES-2100 | AP018178.1:1190855-1192015 |
| *Calothrix* sp. NIES-4071 | AP018255.1:772595-773998 |
| *Calothrix* sp. NIES-4071 | AP018255.1:7573114-7574265 |
| *Calothrix* sp. NIES-4105 | AP018290.1:7570946-7572097 |
| *Calothrix* sp. NIES-4105 | AP018290.1:c773967-772564 |
| *Crinalium epipsammum* PCC 9333 | CP003620.1:c2520624-2519380 |
| *Cyanobacterium aponinum* PCC 10605 | CP003947.1:1114945-1116162 |
| *Cyanothece* sp. PCC 7822 plasmid Cy782201 | CP002199.1:360048-361412 |
| *Cylindrospermum licheniforme* UTEX B 2014 | KX682397.1:2961-4376 |
| *Cylindrospermum* sp. NIES-4074 | AP018269.1:2484675-2486075 |
| *Cylindrospermum stagnale* PCC 7417 | CP003642.1:c2217555-2216140 |
| *Cylindrospermum stagnale* PCC 7417 | CP003642.1:c2308133-2306952 |
| *Cylindrospermum stagnale* PCC 7417 plasmid pCYLST.01 | CP003643.1:56891-58261 |
| *Fischerella* sp. NIES-4106 | AP018298.1:1441529-1442746 |
| *Fremyella diplosiphon* NIES-3275 | AP018233.1:1201630-1202805 |
| *Geitlerinema* sp. PCC 7407 | CP003591.1:c2644190-2642961 |
| *Microcoleus* sp. PCC 7113 | CP003630.1:c2944932-2943757 |
| *Moorea bouillonii* PNG5-198 | KP715425.1:20564-21952 |
| *Moorea bouillonii* PNG5-198 | KP715425.1:22089-23501 |
| *Moorea producens* JHB | CP017708.1:c2514441-2513215 |
| *Moorea producens* PAL-8-15-08-1 | CP017599.1:c2474933-2473707 |
| *Nodularia* sp. HBU26 | KY594676.1:18618-19922 |
| *Nostoc carneum* NIES-2107 | AP018180.1:4766806-4767993 |
| *Nostoc commune* HK-02 | AP018326.1:6163477-6164874 |
| *Nostoc flagelliforme* CCNUN1 | CP024785.1:1135915-1137312 |
| *Nostoc punctiforme* PCC 73102 | CP001037.1:4184899-4186269 |
| *Nostoc punctiforme* PCC 73102, | CP001037.1:c5828161-5826956 |
| *Nostoc* sp. CAVN2 | KT826756.1:3912-5327 |
| *Nostoc* sp. CCAP 1453/38 | KP143720.1:21035-22441 |
| *Nostoc* sp. CCAP 1453/38 | KP143720.1:22490-23818 |
| *Nostoc* sp. CENA543 | CP023278.1:c1270872-1269652 |
| *Nostoc* sp. '*Lobaria pulmonaria'* (5183) | CP026692.1:3554655-3555860 |
| *Nostoc* sp. NIES-4103 | AP018288.1:c2213285-2211918 |
| *Nostoc* sp. NIES-4103 | AP018288.1:c3294496-3293132 |
| *Nostoc* sp. PCC 7524 | CP003552.1:c5619284-5618085 |
| *Nostoc* sp. 'Peltigera membranacea cyanobiont' N6 | CP026681.1:c3280480-3279275 |
| *Nostoc* sp. UIC 10110 | KY379971.1:817-2223 |

**Table S1.** (continued)

| | |
|---|---|
| *Nostocales* cyanobacterium HT-58-2 | CP019636.1:1098629-1099993 |
| *Oscillatoria acuminata* PCC 6304 | CP003607.1:c70363-69134 |
| *Pleurocapsa* sp. PCC 7327 | CP003590.1:298214-299635 |
| *Rivularia* sp. PCC 7116 | CP003549.1:c7519531-7518356 |
| *Stanieria* sp. NIES-3757 | AP017376.1:c59518-58163 |
| *Streptomyces thioluteus* HKI-22 | AJ575648.1:4858-5868 |
| *Synechocystis salina* LEGE 06099 | KX083339.1:12314-13522 |
| *Synechocystis salina* LEGE 06155 | KR059027.1:13578-14783 |
| *Synechocystis* sp. IPPAS B-1465 | CP028094.1:c2118162-2116924 |
| *Synechocystis* sp. PCC 6714 | CP007542.1:c2270478-2269219 |
| *Synechocystis* sp. PCC 6803 | CP003265.1:c2114589-2113351 |
| *Tolypothrix tenuis* PCC 7101 | AP018248.1:c3613364-3612189 |

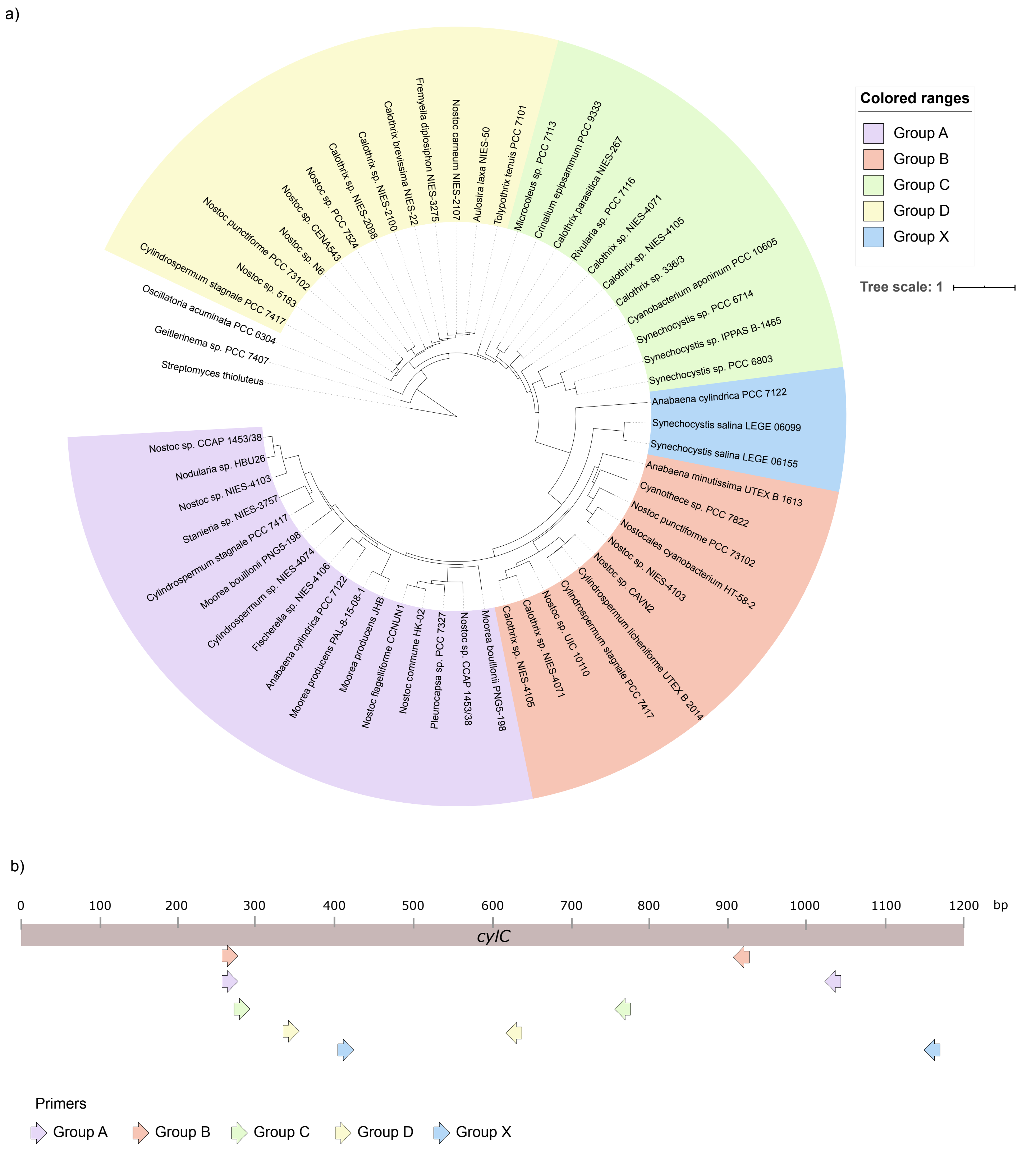


**Figure S1.** (a) Phylogenetic tree (FastTree GTR with a rate of 100) of *cylC* homologs highlighted according to the groups selected for degenerate primer design. (b) Schematic representation of the different pairs of degenerate primers

Group of primers A

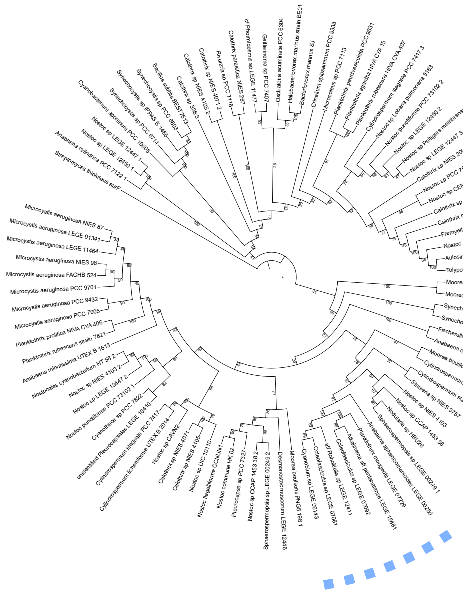

Group of primers B

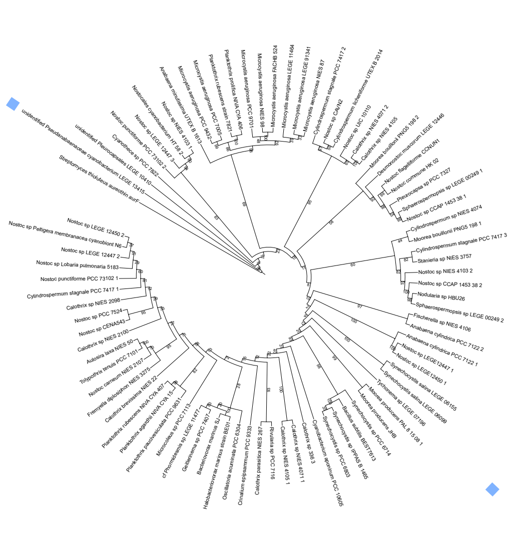

Group of primers C

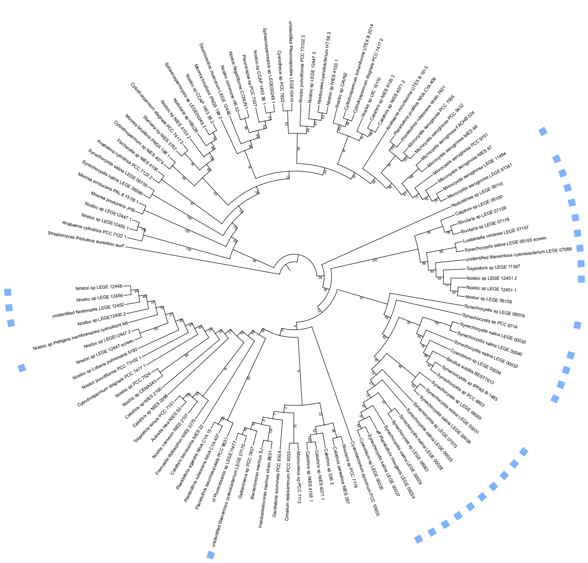

Group of primers D

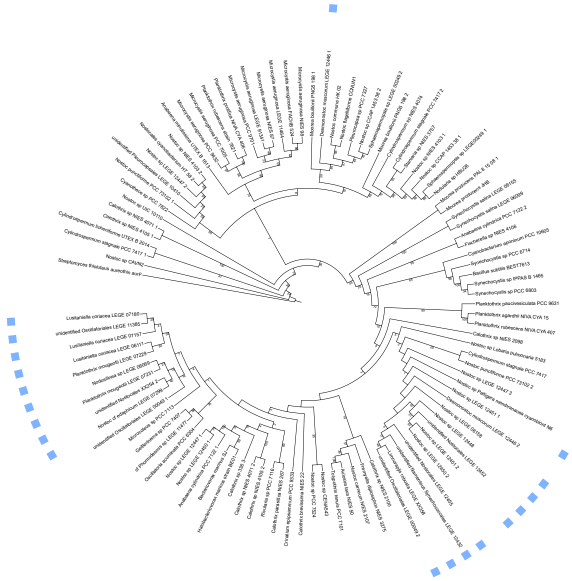

Group of primers X

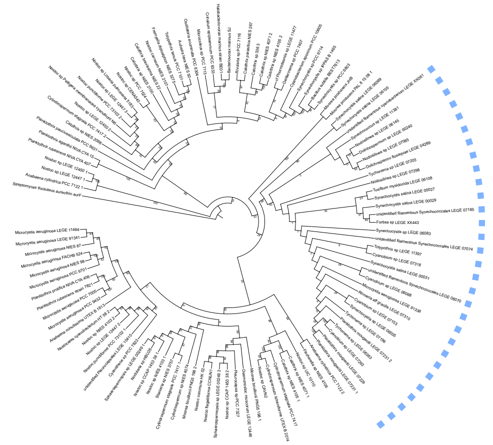

**Figure S2.** PCR-based detection of *cylC* homologs in the LEGEcc culture collection. Five pairs of primers were designed based on conserved regions identified in the *cylC* gene. Each primer pair was used in a PCR screen of the gDNA obtained from diverse strains (n = 326) of the LEGEcc culture collection. The resulting amplicons were cloned and sequenced. Sequences for each primer pair were aligned with the corresponding regions of *cylC* genes found in the NCBI reference genomes (cyanobacteria only) and those from LEGEcc strains' genomes. Shown are the resulting cladograms (RaxML, 1000 replicates) for each primer pair used in the screening. Blue squares indicate sequences obtained from the PCR screen.

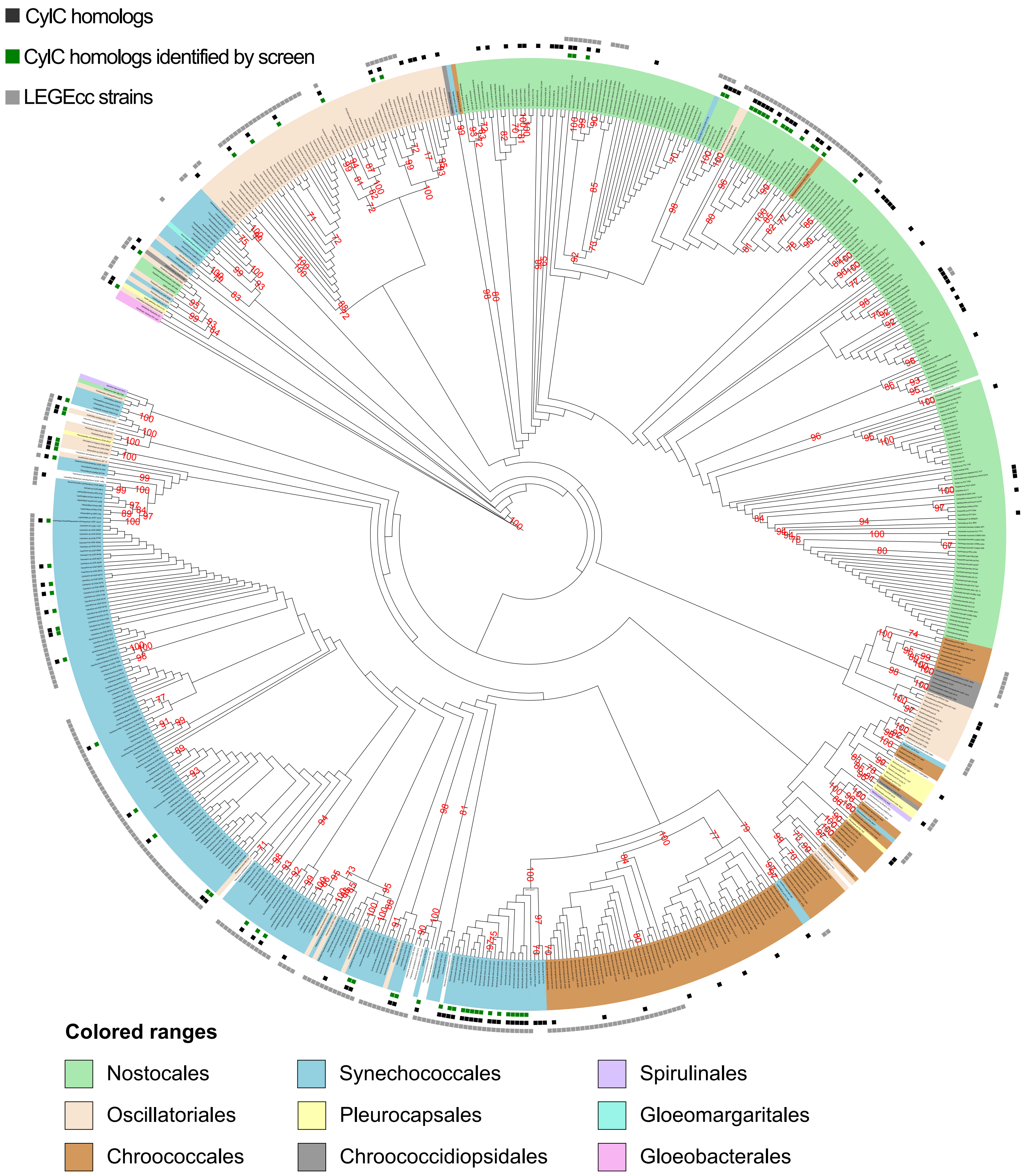


**Figure S3.** RaxML cladogram (1000 replicates) of the 16S rRNA gene of LEGEcc strains (grey squares) and from cyanobacterial strains with NCBI-deposited reference genomes, screened in this study. Taxonomy is presented at the order level (colored ranges). Strains whose genomes encode CylC homologs are denoted by black squares. Green squares indicate that at least one CylC homolog was detected by PCR-screening and verified by retrieving the sequence of the corresponding amplicon through cloning followed by Sanger sequencing. The cladogram topology is the same as shown in Figure 3 of the main manuscript, but here bootstrap values (equal or above 0.7) are shown.

**Table S2.** GenBank or RefSeq assembly acession number and LEGEcc genome used for CORASON analysis.

| Enzyme | Reference genome | Number of genomes/Clusters tested in CORASON | Removed from the alignment | Included manually extracted BGCs from genomes |
|---|---|---|---|---|
| CylC | GCA_000317535.1 | 2170 | GCA_003504865.1, LEGE00239, GCA_000332035.1, GCA_004305995.1, GCA_003206555.1, GCA_000312205.1 | CP011382.1, CP026692.1, NZ_LN887838.1, NZ_LN887838.1, KT826756.1, KP143720.1, NZ_CP023280.1, KY594676.1, NZ_CP028099.1, LEGE00239, LEGE_00249, LEGE06071, LEGE06083, LEGE06083, LEGE_06099, LEGE_06155, LEGE07170, LEGE11397, LEGE_11477, LEGE11479, LEGE11480, LEGE12450, LEGE12450, , NZ_CP026681.1, AP018172.1, AP018178.1, AP018180.1, AP018207.1, AP018227.1, AP018233.1, AP017375.1, AP018307.1, NZ_AVFS00000000.1, NZ_AVFV00000000.1, NZ_AVFV00000000.1, NZ_KE734719.1, NC_019776.1", NC_019693.1, CP007542.1, CP003265.1, NZ_AQPY00000000.1, NZ_AP018248.1, CP003630.1, CP003549.1, CP003591.1, CP003642.1, CP003552.1, CP003620.1, NZ_ALVX00000000.1, NZ_ALVX00000000.1, NZ_HE972553.1, NZ_CZCS00000000.2, KY379971.1, MH325199.1, KX682397.1 |
| PrnA | GCA_005518235.1 | 2114 | GCA_002025445.1, GCA_002025445.1 | - |
| CurA | GCA_001942475.1 | 2116 | - | - |
| mcnD | GCA_000312245.1 | 2114 | LEGE 17501, LEGE 91341 | - |
| hs_bmp5 | GCA_001942475.1 | 2114 | GCA_005518205.1, GCA_005518235.1 | - |
| WelO5 | GCA_000447295.1 | 2116 | - | - |

**Table S3.** BLASTp search of CylC homologs against *Aliterella sp.*, *Chroococcidiopsis sp.* and *Gloeobacter sp.*

| Input | CylC | BrtJ | ColE | ColD | NocO | NocN | Mic |
|---|---|---|---|---|---|---|---|
| **Strain/ Accession Number** | ***Aliterella atlantica*/ WP_045054787.1** | | | | | | |
| **Coverage** | 94% | 95% | 93% | 96% | 95% | 97% | 98% |
| **E-value** | 3,00E-147 | 4,00E-67 | 2,00E-153 | 7,00E-176 | 8,00E-124 | 0.0 | 7,00E-148 |
| **Percent identity** | 47.33% | 41.70% | 49.32% | 51.72% | 43.57% | 69.52% | 46.96% |
| **Strain/ Accession Number** | ***Chroococcidiopsis sp. TS-821*/WP_104546385.1** | | | | | | |
| **Coverage** | 98% | 99% | 93% | 99% | 95% | 99% | 94% |
| **E-value** | 4,00E-155 | 9,00E-68 | 8,00E-159 | 7,00E-175 | 1,00E-121 | 0.0 | 6,00E-150 |
| **Percent identity** | 48.65% | 42.98% | 51.93% | 50.85% | 43.15% | 68.66% | 49.22% |
| **Strain/ Accession Number** | ***Gloeobacter violaceus SpSt-379*/HGZ84776.1** | | | | | | |
| **Coverage** | 94% | 94% | 95% | 95% | 100% | 94% | 95% |
| **E-value** | 1,00E-57 | 1E-109 | 5,00E-92 | 1,00E-98 | 3,00E-87 | 3,00E-100 | 5,00E-62 |
| **Percent identity** | 42.79% | 43.98% | 36.00% | 37.64% | 34.39% | 37.16% | 46.73% |

***Calothrix brevissima* NIES-22 DNA**
AP018207.1

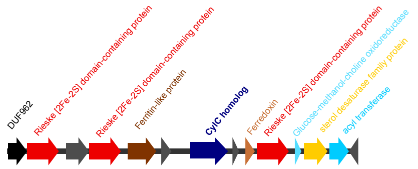


***Tolypothrix tenuis* PCC 7101**
AP018248.1

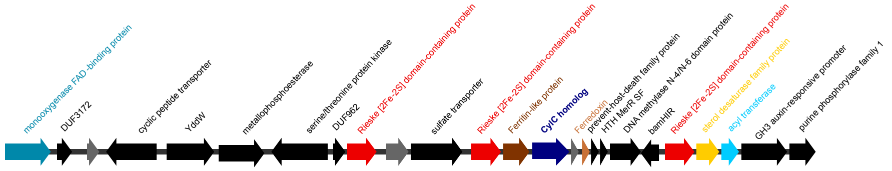


***Nostoc punctiforme* PCC 73102**
CP001037.1
Cluster 2

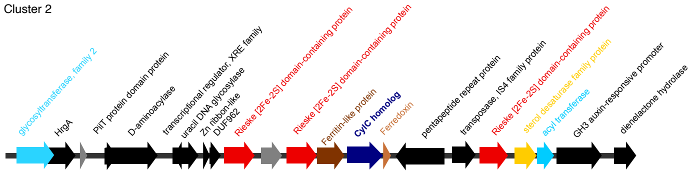


***Aulosira laxa* NIES-50**
AP018307.1

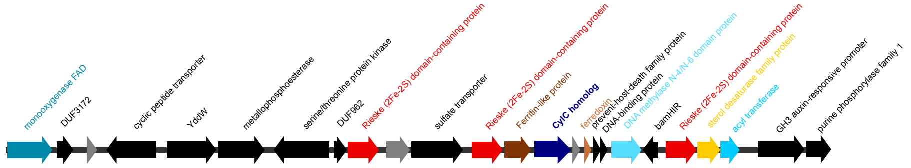


***Calothrix* sp. NIES-4071**
AP018255.1
Cluster 1

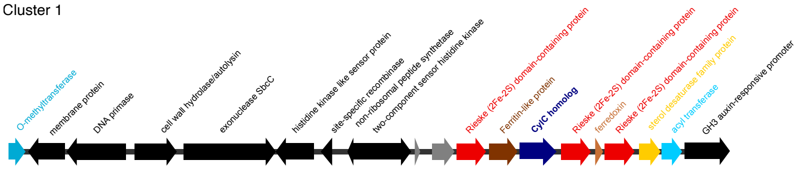


***Calothrix* sp. NIES-4105**

AP018290.1
Cluster 1

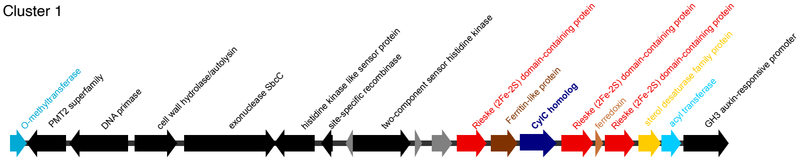


***Calothrix parasitica* NIES-267**
AP018227.1

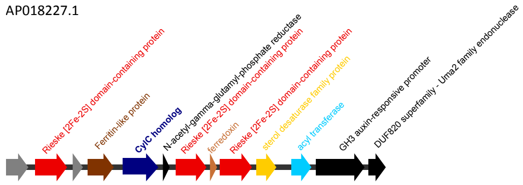

***Calothrix* sp. NIES-2100**

AP018178.1

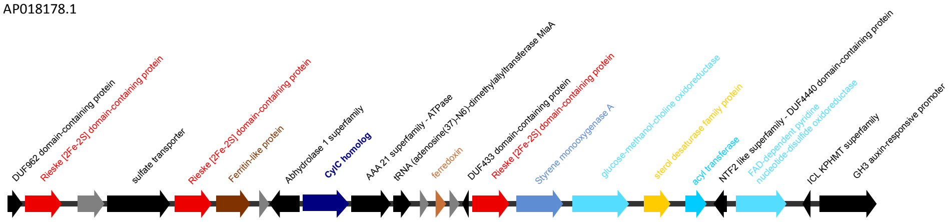


***Geitlerinema* sp. PCC 7407**

CP003591.1

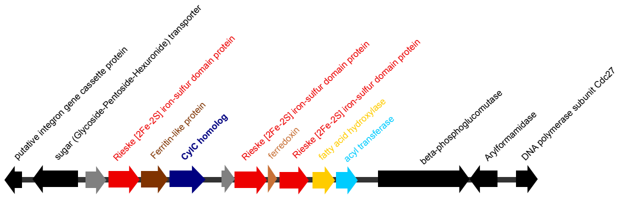


***Nostoc* sp. CENA543**

CP023278.1

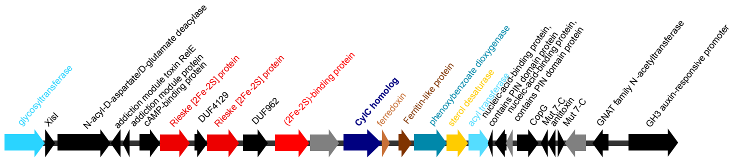


***Cyanobacterium aponinum* PCC 10605**

CP003947.1

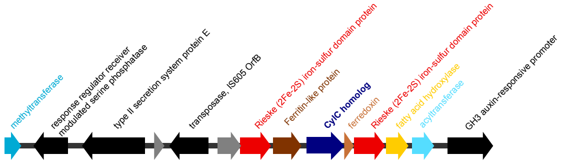


***Calothrix* sp. NIES-2098 DNA**

AP018172.1

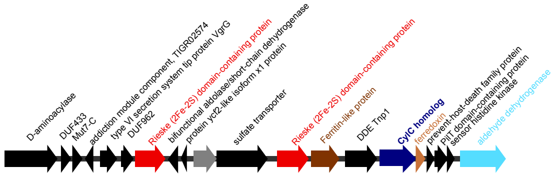


***Nostoc* sp. 'Peltigera membranacea cyanobiont' N6**

CP026681.1

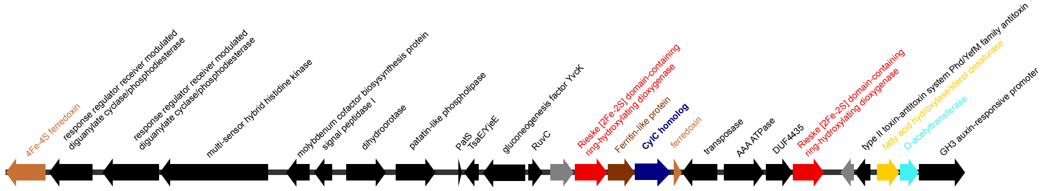


***Nostoc carneum* NIES-2107**

AP018180.1

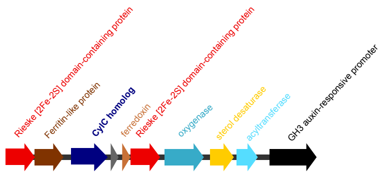

**Fremyella diplosiphon NIES-3275**
AP018233.1

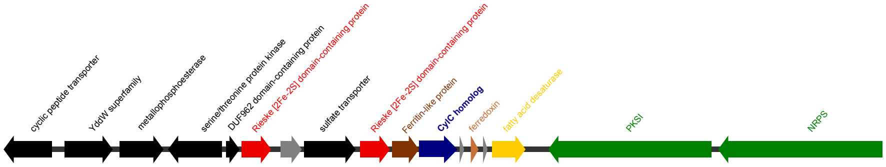


**cf. *Phormidesmis sp.* LEGE 11477**

**Cluster 1**

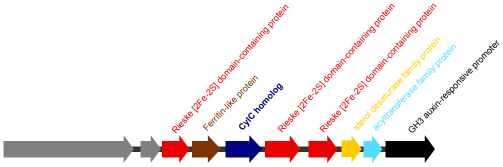


***Nostoc sp.* LEGE 12447**

**Cluster1**

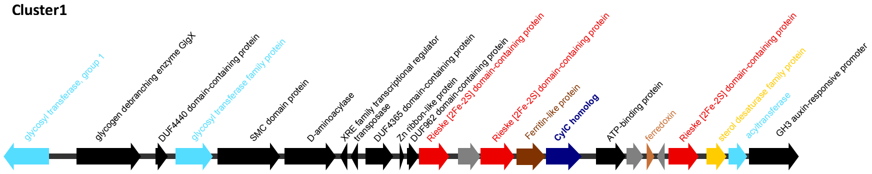


***Nostoc sp.* LEGE 12450**

**Cluster1**

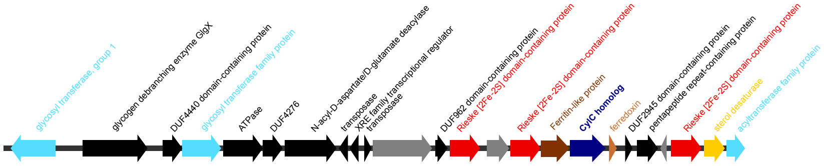


***Planktothrix rubescens* strain 7821**
**Cluster 1**

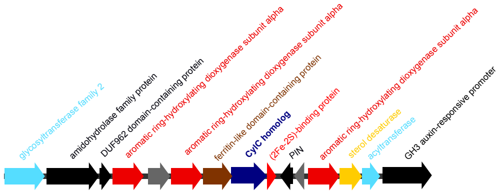


***Microcoleus* sp. PCC 7113**
CP003630.1

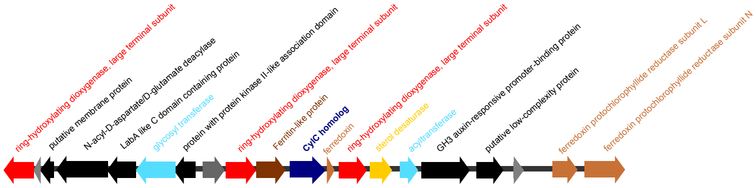


***Cylindrospermum stagnale* PCC 7417**
CP003642.1
Cluster2

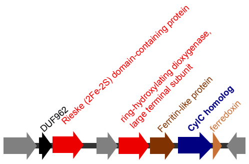

***Oscillatoria acuminata* PCC 6304**
CP003607.1

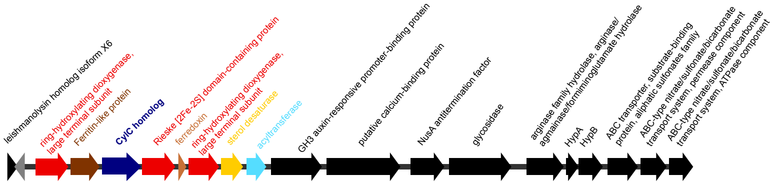


***Rivularia* sp. PCC 7116**
CP003549.1

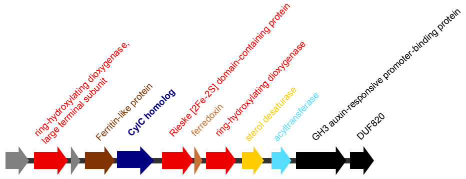


***Crinalium epipsammum* PCC 9333**
CP003620.1

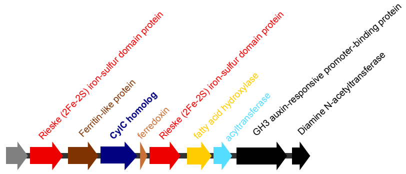


***Calothrix* sp. 336/3**
CP011382.1

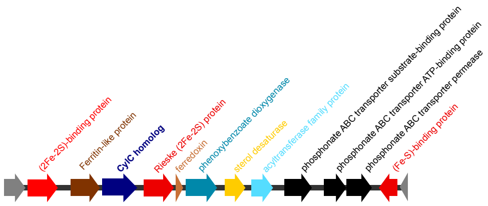


***Nostoc* sp. 'Lobaria pulmonaria (5183) cyanobiont' strain 5183**
CP026692.1

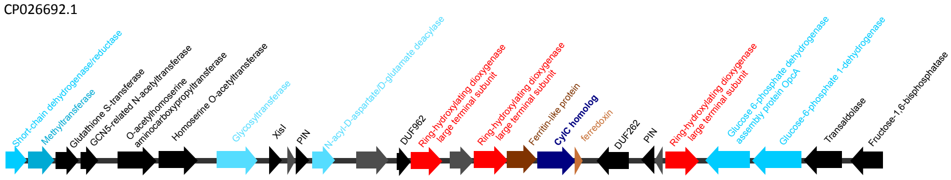


***Synechocystis* sp. LEGE 06083**
**Cluster 1**

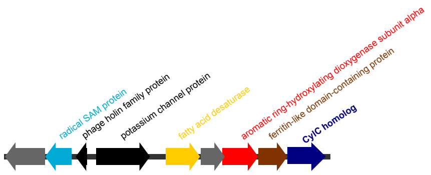


***Leptolyngbya saxicola* LEGE 07170**

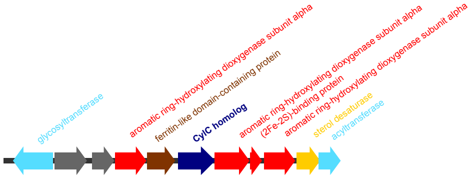

***Tolypothrix* sp. LEGE 11397**

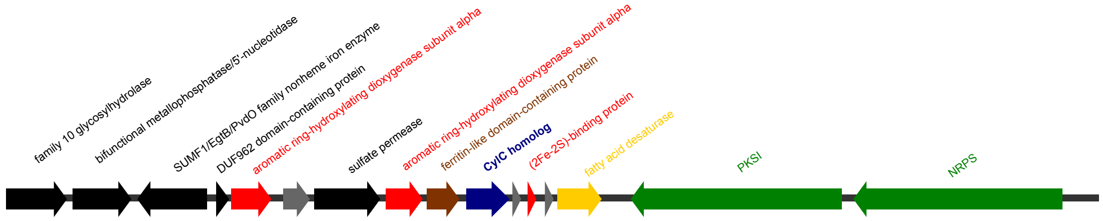


***Leptolyngbya cf. ectocarpi* LEGE 11479**

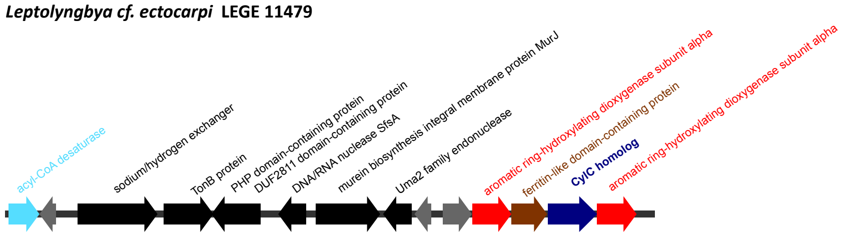


**unidentified filamentous cyanobacterium LEGE 11480**

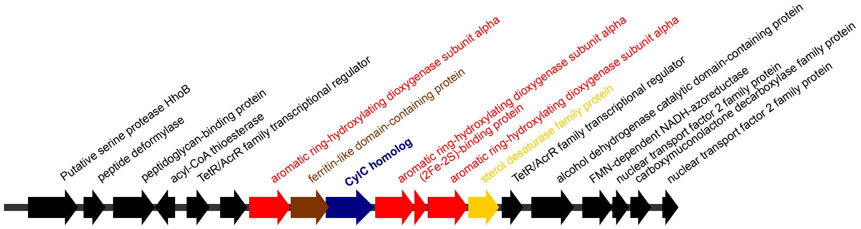


***Planktothrix agardhii* NIVA CYA 15**

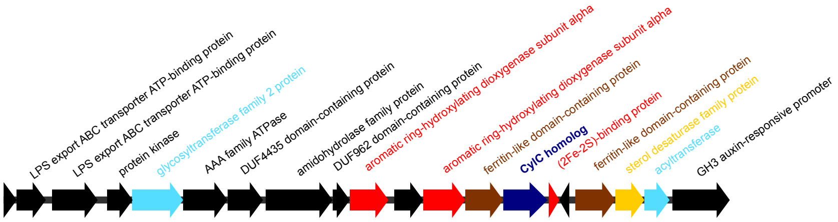


***Planktothrix rubescens* NIVA CYA 406**
**Cluster 1**

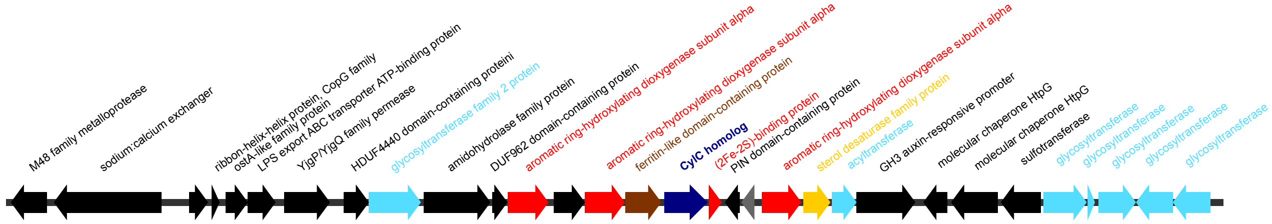


***Planktothrix rubescens* NIVA CYA 407**

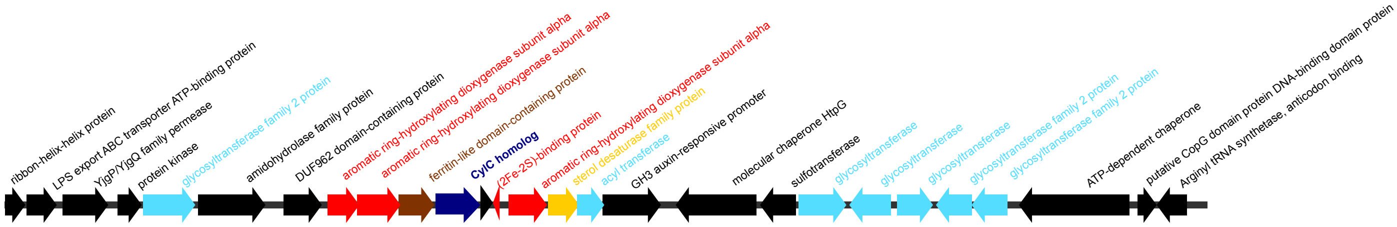

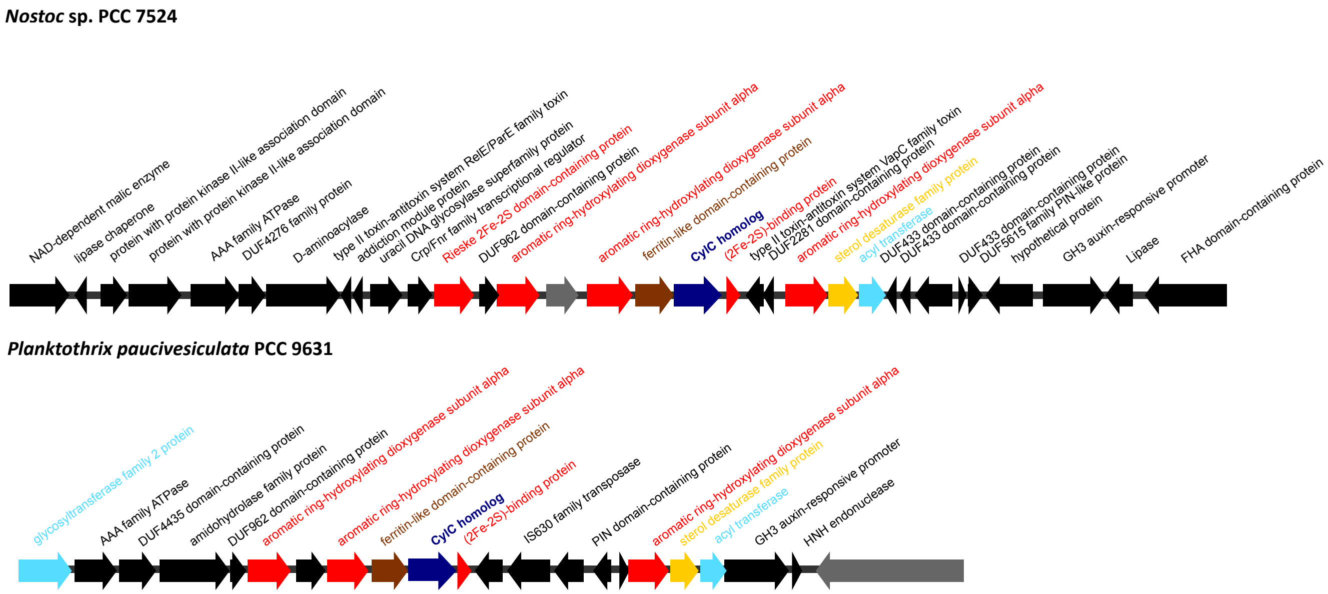


**Figure S4.** Rieske-containing biosynthetic gene clusters encoding CylC homolog(s).

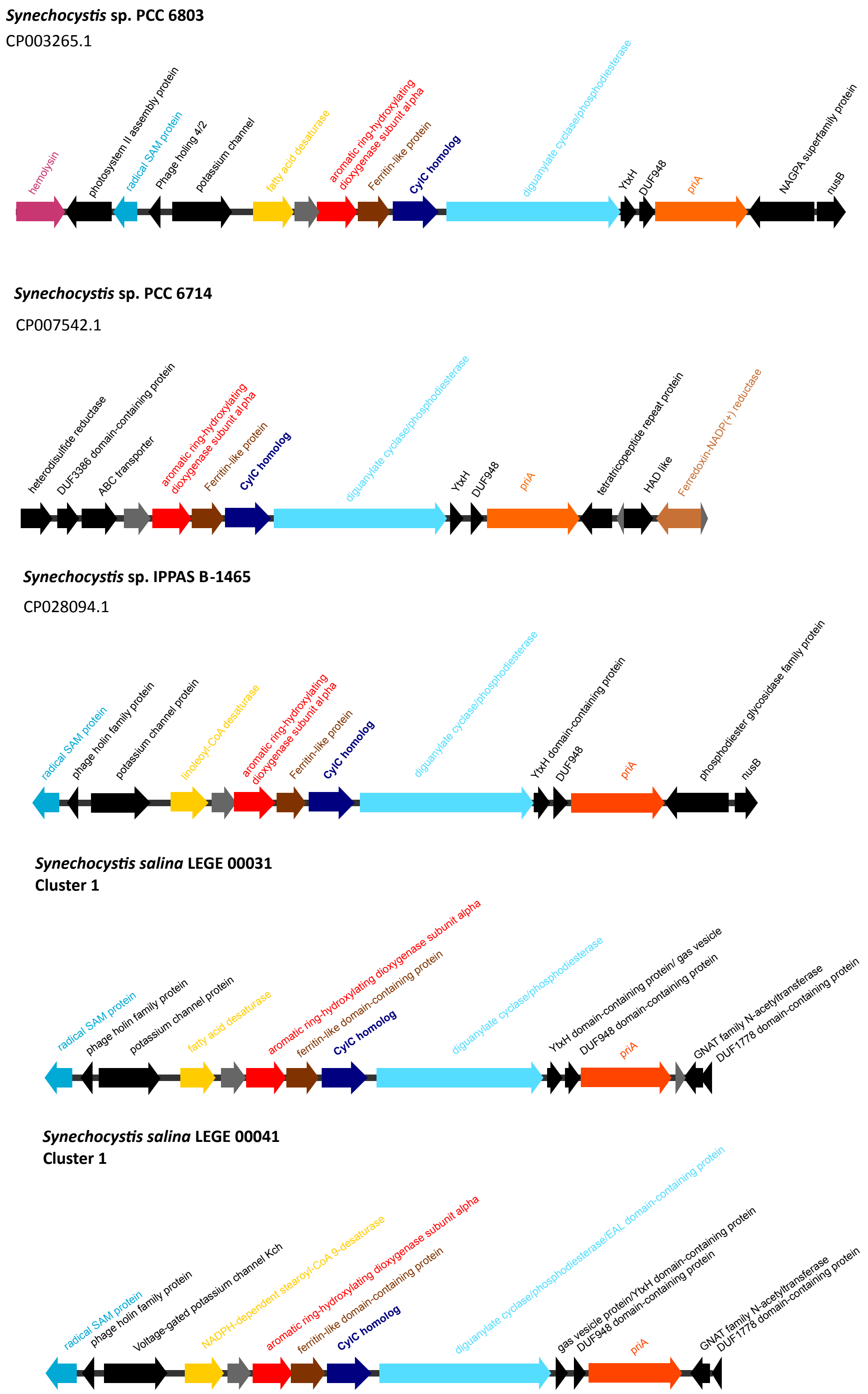


**Figure S5.** PriA-containing biosynthetic gene clusters encoding CylC homolog(s).

***Stanieria* sp. NIES-3757**

AP017376.1

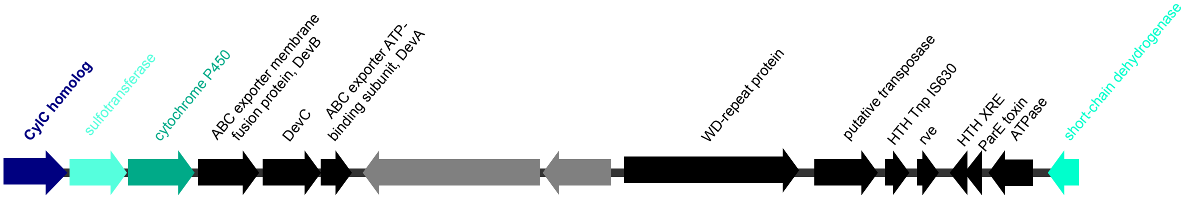


**Figure S6. C**ytochrome P450/sulfotransferase-containing biosynthetic gene cluster encoding a CylC homolog.

***Nostoc* sp. CCAP 1453/38**

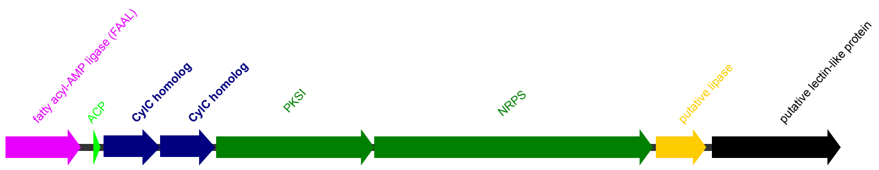


***Nodularia* sp. HBU26**
KY594676.1

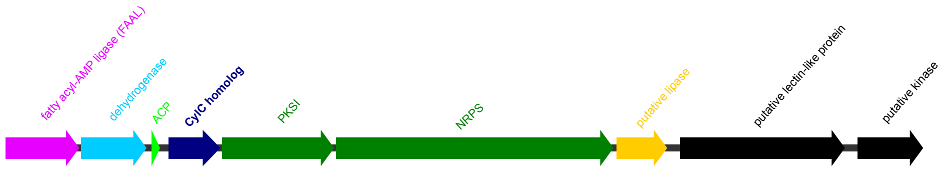


***Sphaerospermopsis* sp. LEGE00249**

*chlorosphaerolactylates (cly)*

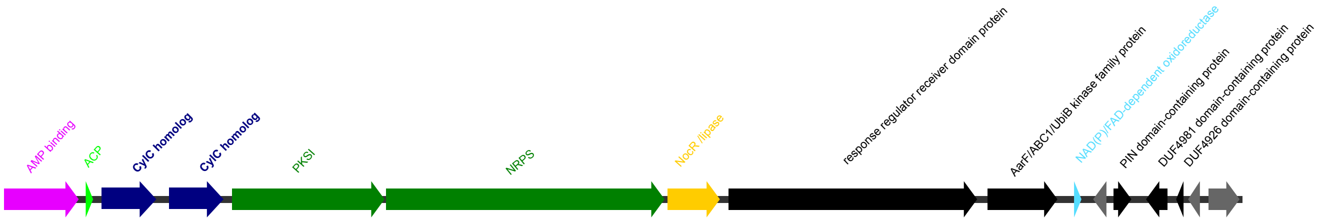


***Anabaena aphanizomenoides* LEGE 00250**

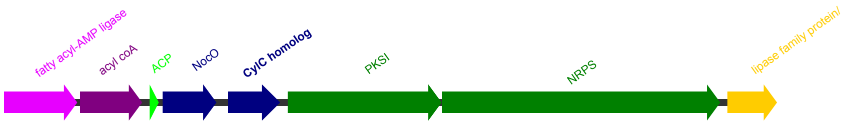

***Nodularia* sp. LEGE 06071**

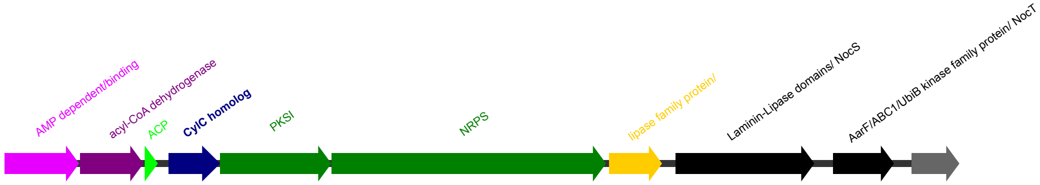


***Moorea bouillonii* PNG5-198**

KP715425.1 *columbamides (col)*

**Cluster1**

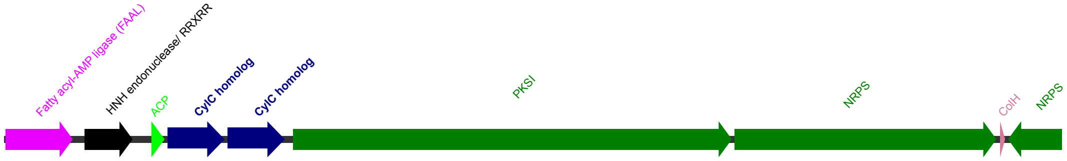


***Microcystis aeruginosa* 91341**

**Cluster1**

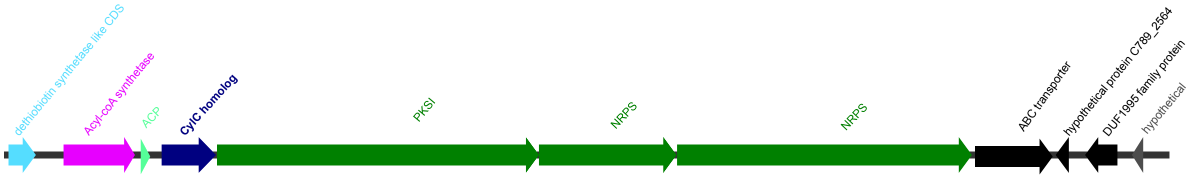

***Microcystis aeruginosa* PCC 9432**

*microginin (mic)*

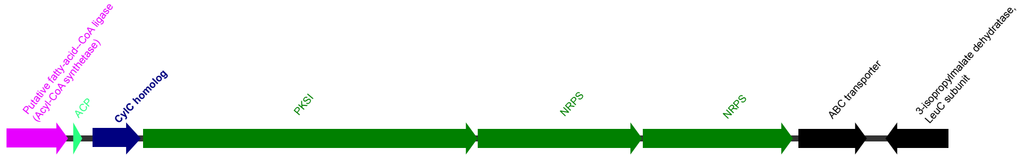


***Microcystis aeruginosa* FACHB-524**

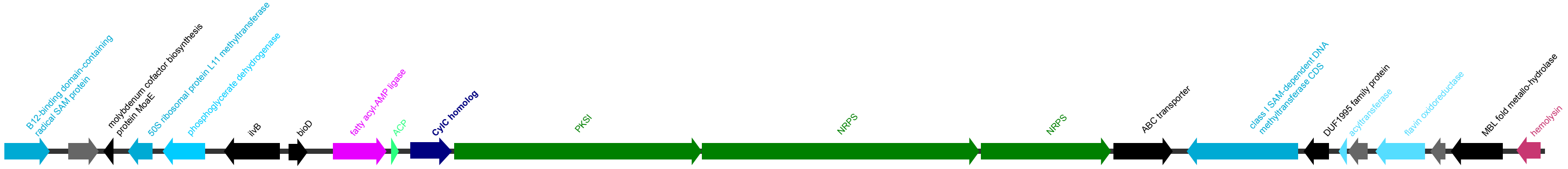


***Planktothrix rubescens* strain 7821**

**Cluster 2**

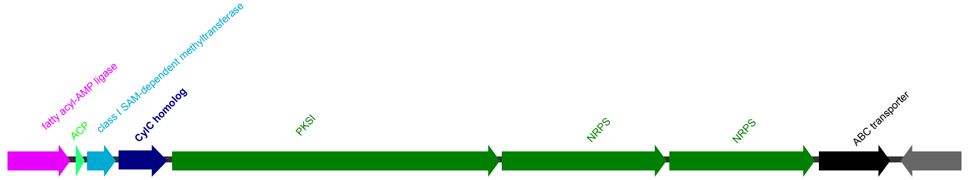

***Microcystis aeruginosa* NIES 87**

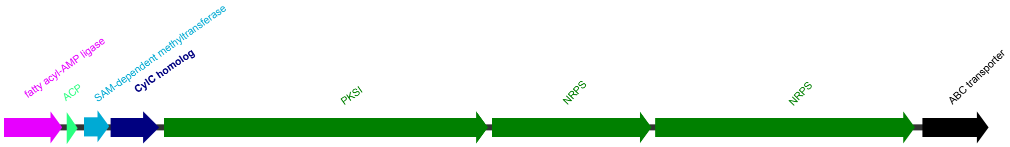


***Microcystis aeruginosa* NIES 98**

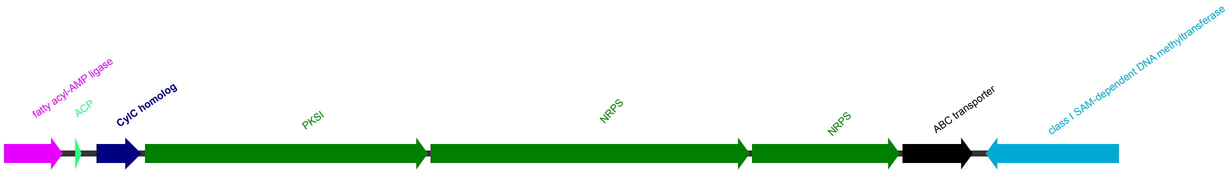


***Planktothrix rubescens* NIVA CYA 406**
**Cluster 2**

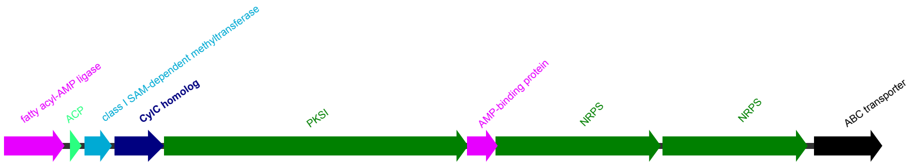


***Microcystis aeruginosa* PCC 7005**

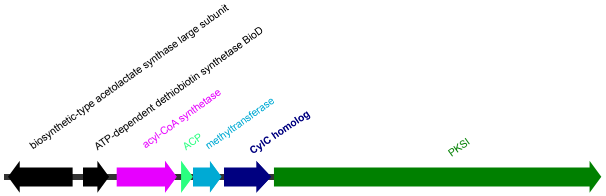

***Cylindrospermum stagnale* PCC 7417 plasmid**

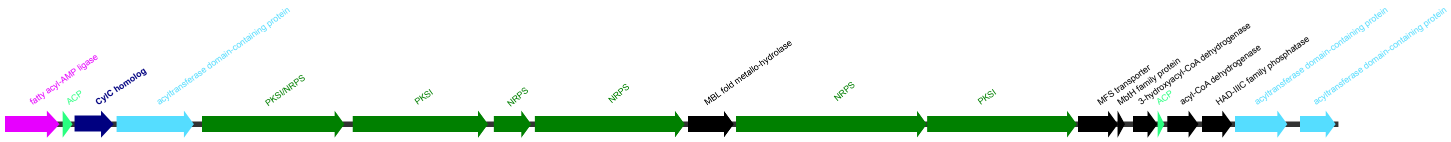


***Fischerella* sp. PCC 9431**
**Cluster2**

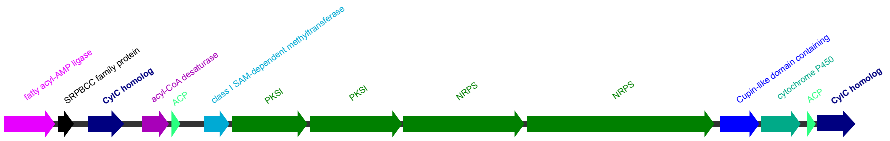


***Nostocales* cyanobacterium HT-58-2**

CP019636

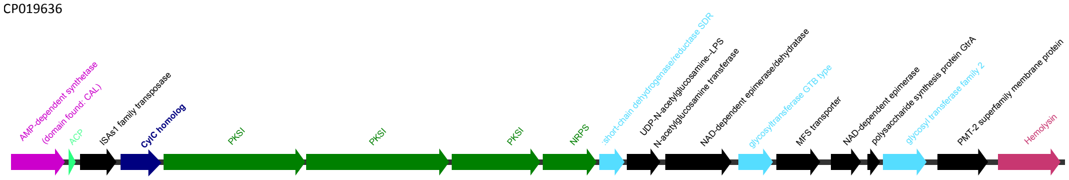


***Anabaena minutissima* UTEX B 1613**

MH325199.1

*minutissamides (puwainaphycin-like, puw)*

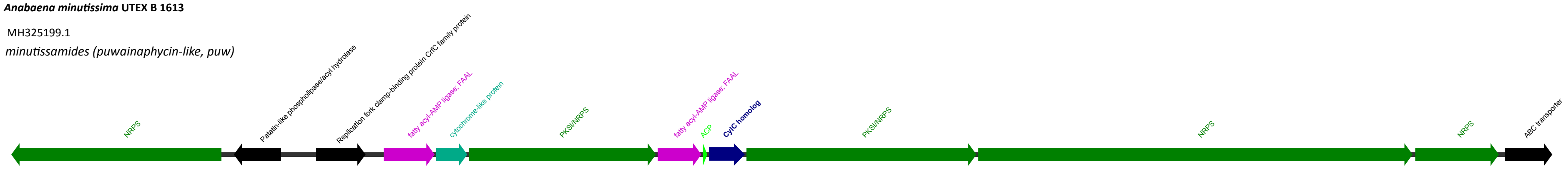

***Pleurocapsa* sp. PCC 7327**

CP003590.1

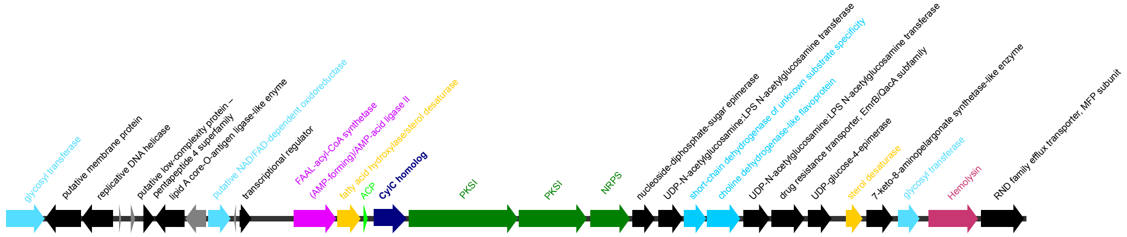


***Desmonostoc muscorum* LEGE 12446**

**Cluster 1**

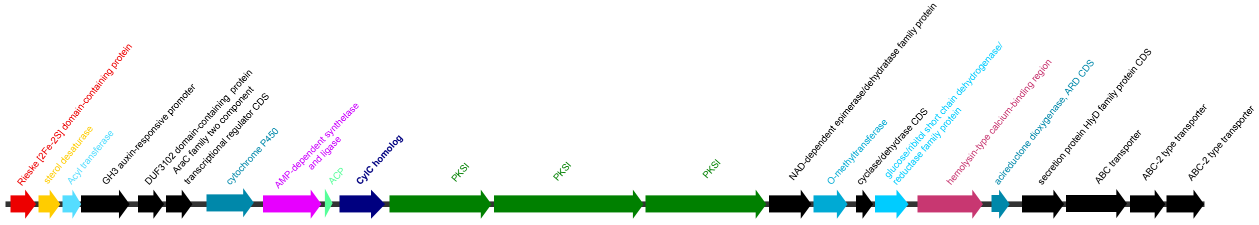


***Nostoc commune* HK-02**

AP018326.1

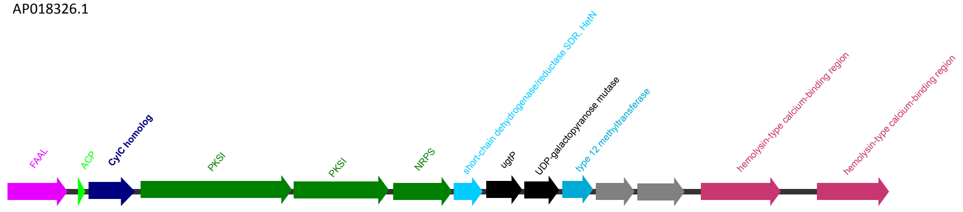

CP001037.1

Cluster1

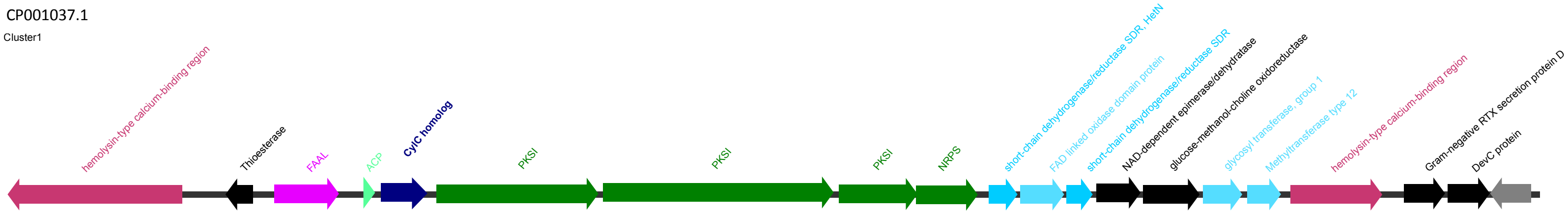


***Nostoc sp.* LEGE 12447**

**Cluster 3**

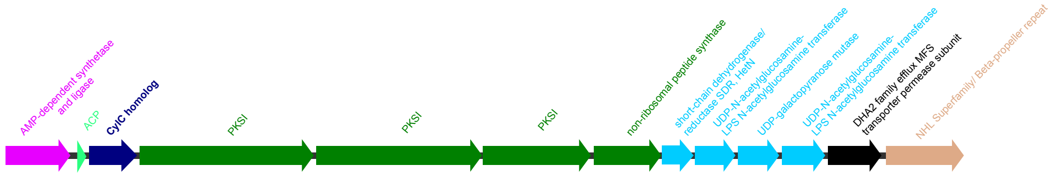


***Nostoc* sp. NIES-4103**

AP018288.1

Cluster 2

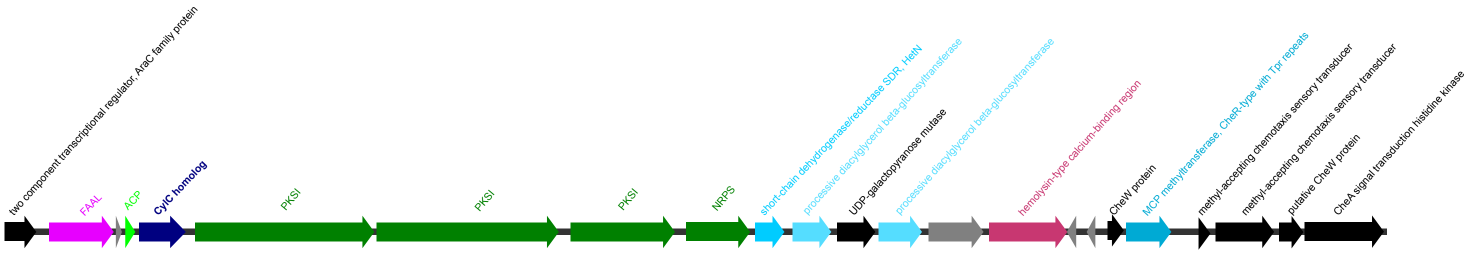


***Cyanothece* sp. PCC 7822**

CP002199.1

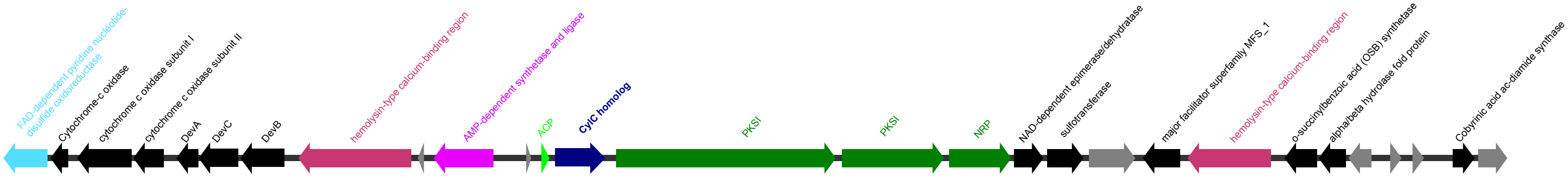

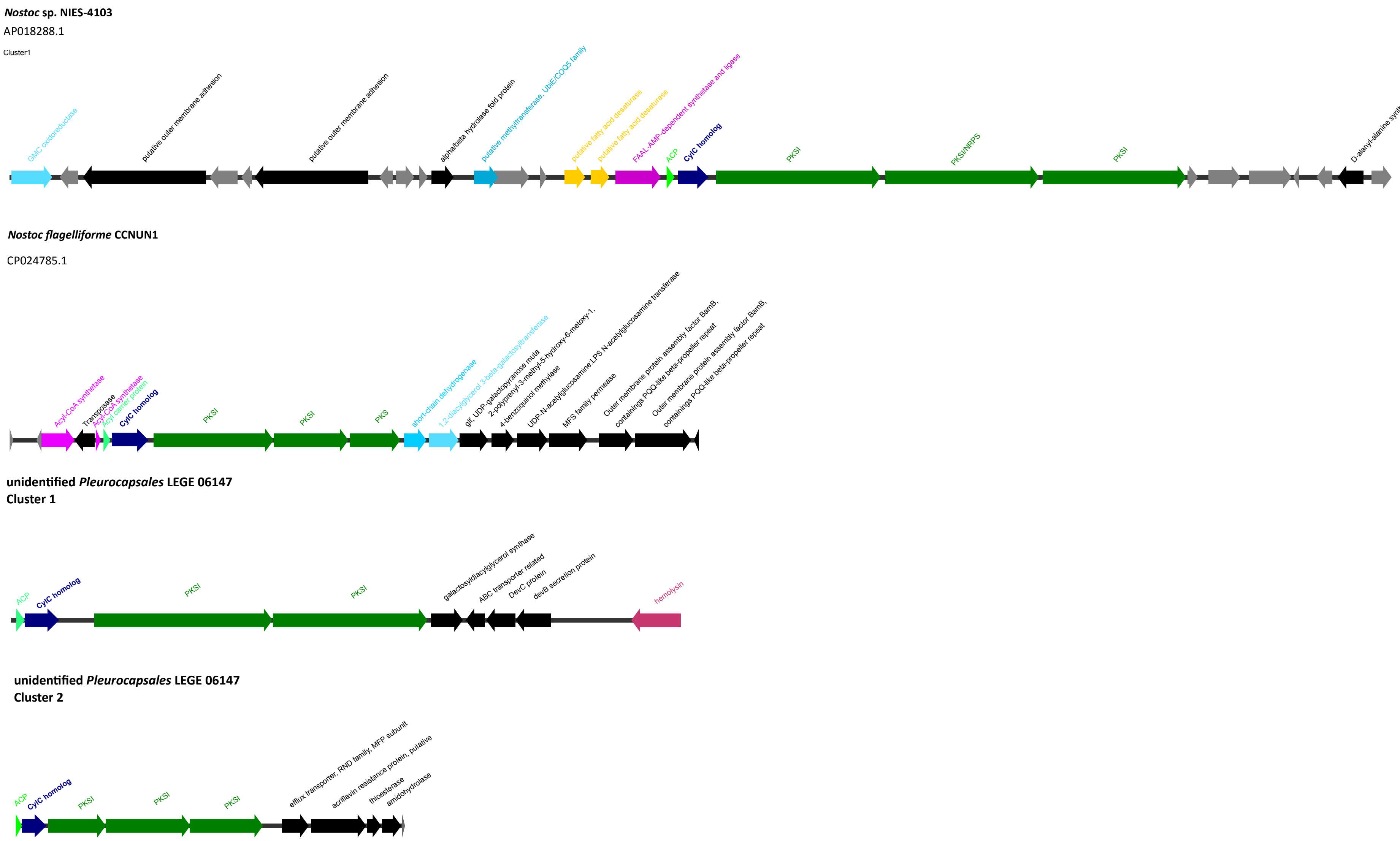


**Figure S7.** Type I PKS (chlorosphaerolactylate/columbamide/microginin/puwainaphycin-like) biosynthetic gene clusters encoding CylC homolog(s).

***Synechocystis salina* LEGE 06155** Bartolosides

KR059027.1

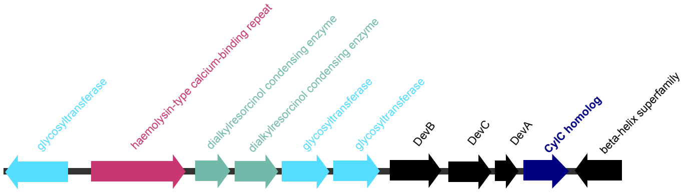


***Synechocystis salina* LEGE 06099** Bartolosides

KX083339.1

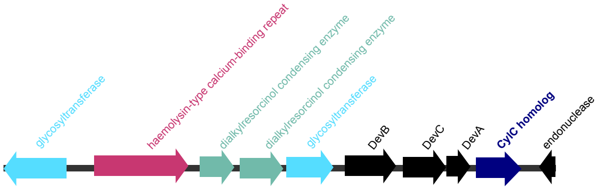


***Synechocystis salina* LEGE 00041**
**Cluster 2**

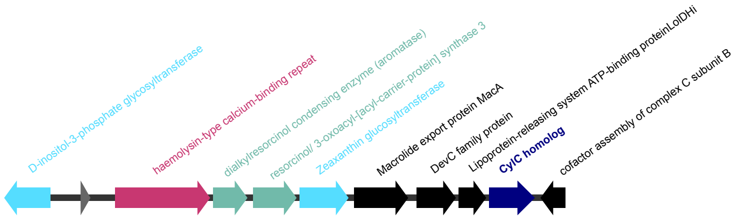


***Fischerella* sp. NIES-4106**

AP018298.1

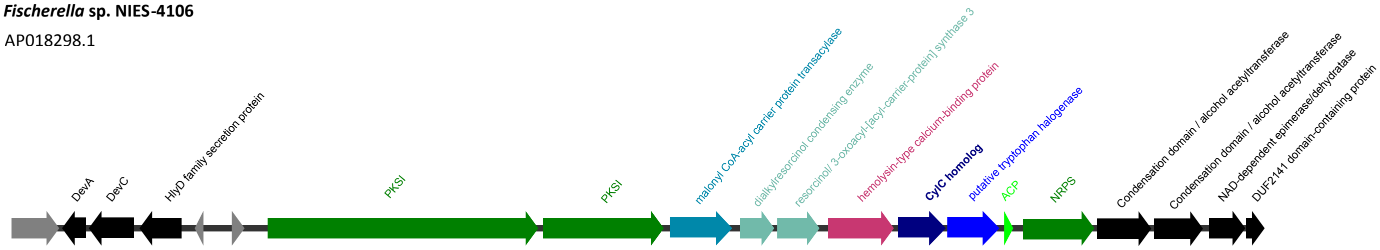

***Anabaena cylindrica* PCC 7122**

CP003659.1 AP018166.1

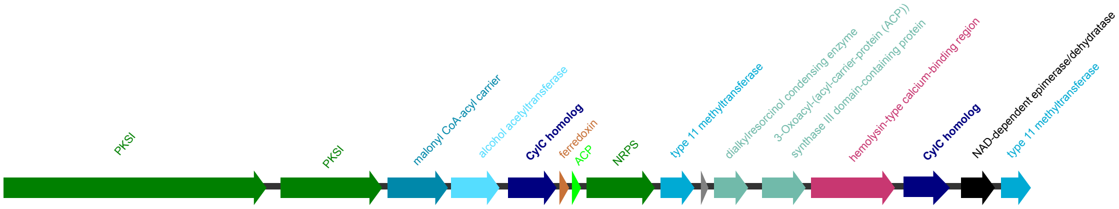


***Synechocystis salina* LEGE 00031**

**Cluster 2**

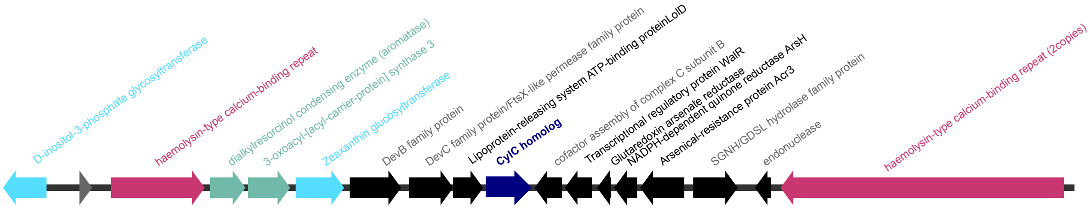


***Synechocystis* sp. LEGE 06083**

**Cluster 2**

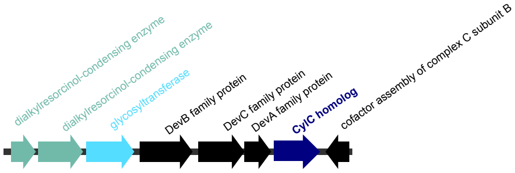


***Fischerella* sp. PCC9431**

**Cluster1**

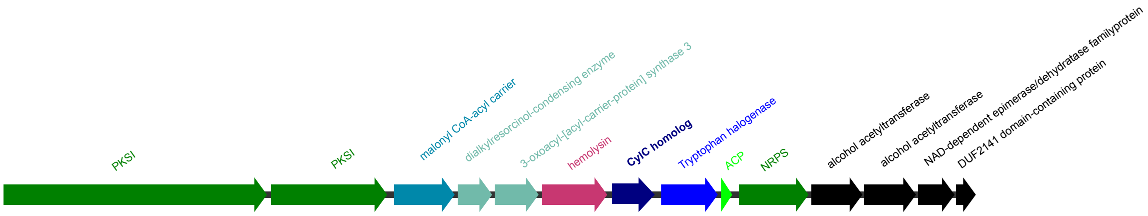


**Figure S8.** Dialkylresorcinol biosynthetic gene clusters encoding CylC homolog(s).

***Cylindrospermum licheniforme* UTEX B 2014** Cylindrocyclophane

KX682397.1

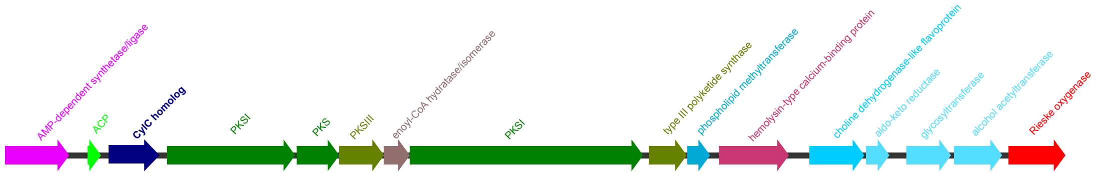


***Cylindrospermum stagnale* PCC 7417**

CP003642.1

Cluster1

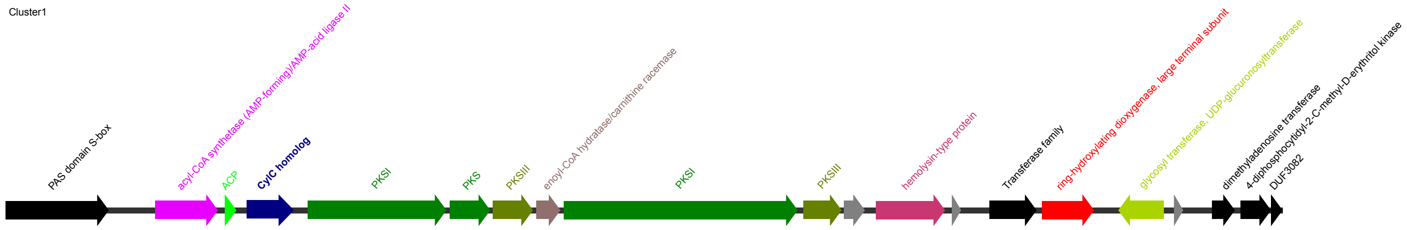


***Nostoc* sp. UIC 10110** Merocyclophane

KY379971.1

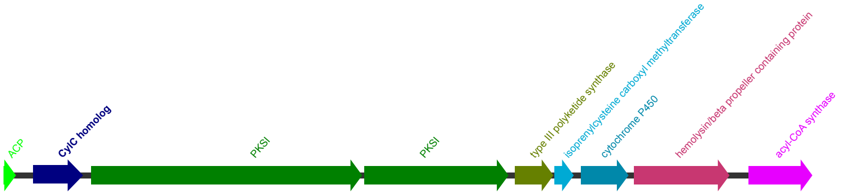


***Nostoc* sp. CAVN2** Carbamidocyclophane

KT826756.1

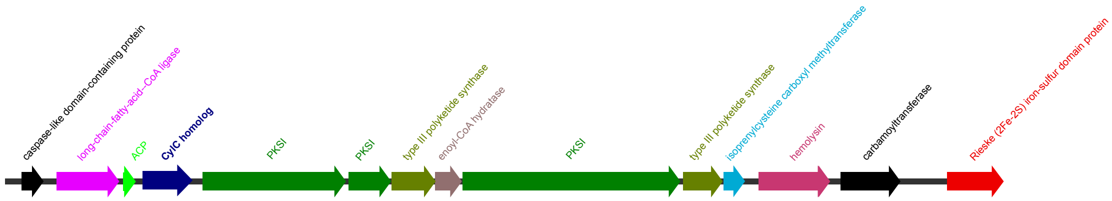

***Cylindrospermum* sp. NIES-4074**

AP018269.1

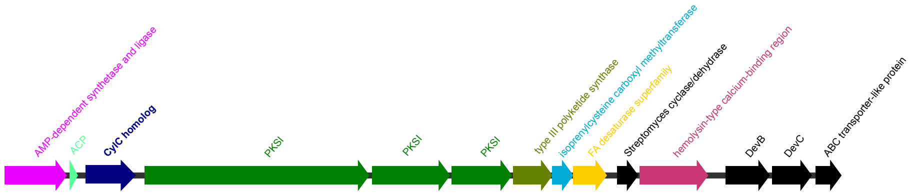


***Calothrix* sp. NIES-4071**

AP018255.1

Cluster2

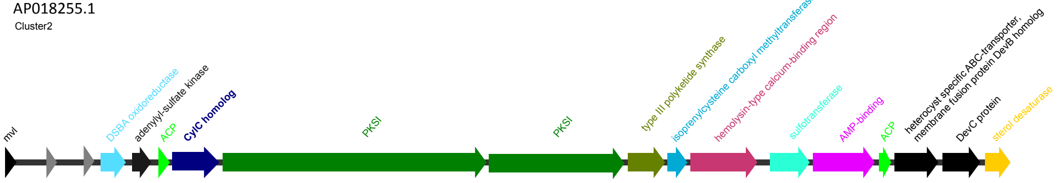


***Calothrix* sp. NIES-4105**

AP018290.1

Cluster2

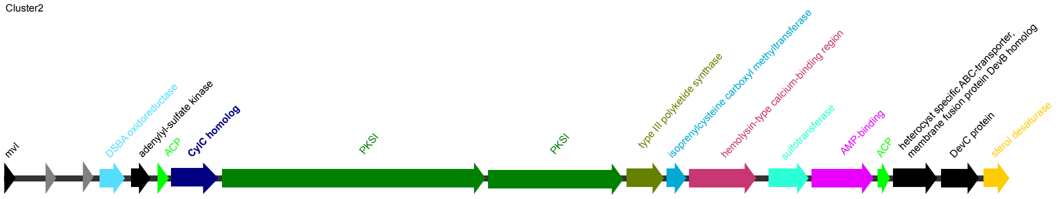


**unidentified *Pleurocapsales* LEGE 10410**

**Cluster 1**

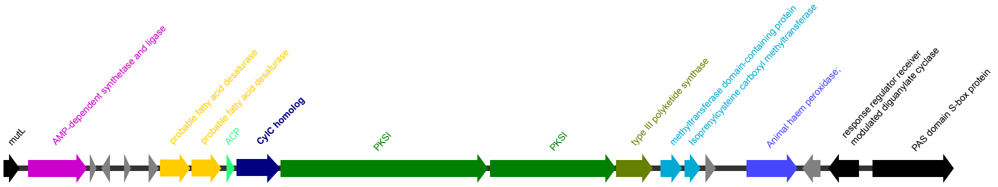


***Hyella patelloides* LEGE 07179**

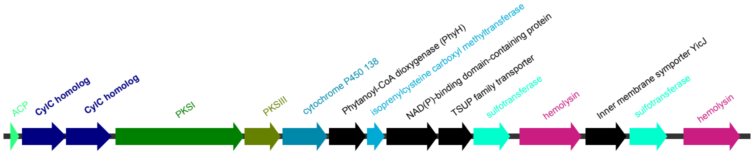

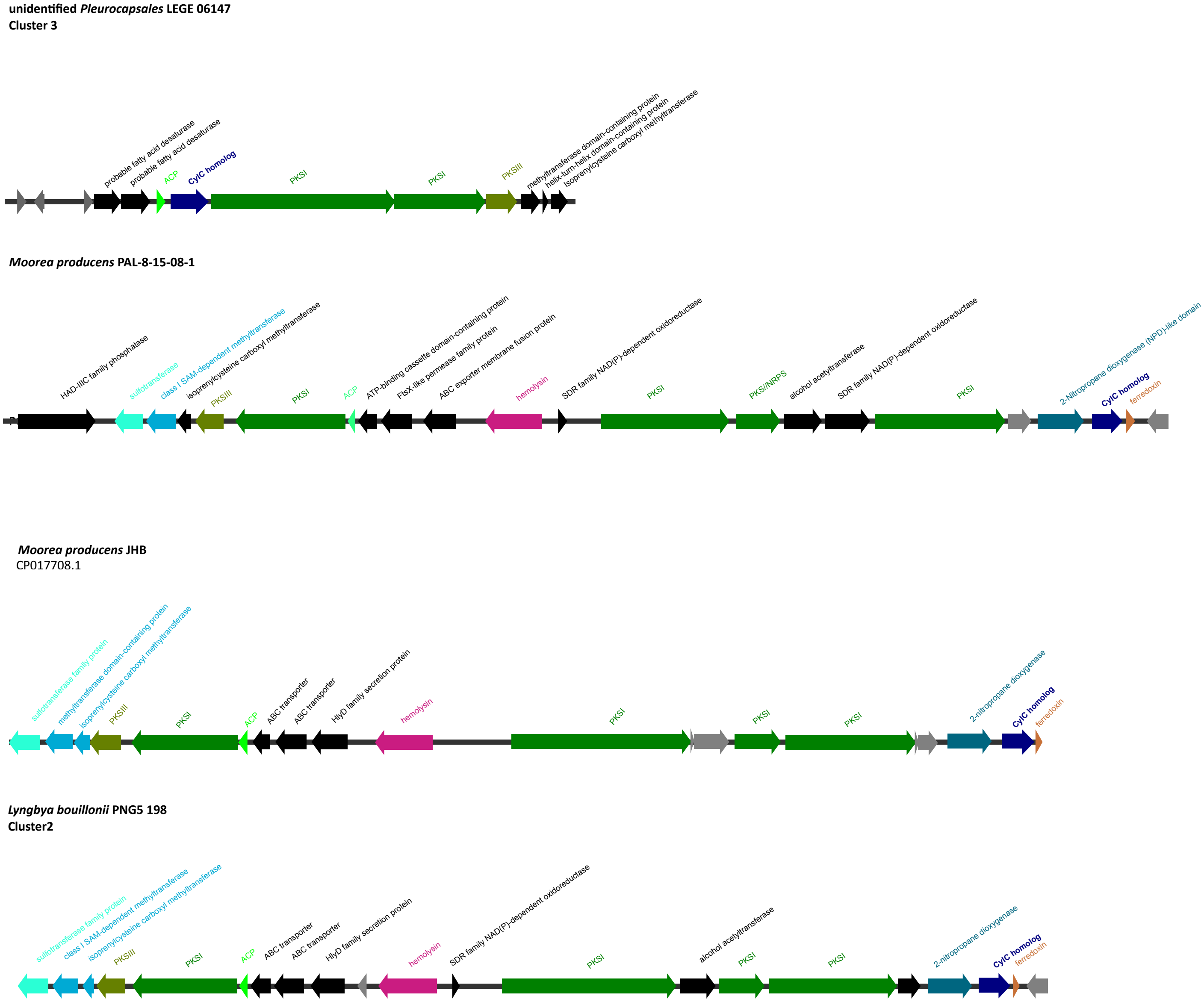


**Figure S9.** Type III PKS biosynthetic gene clusters encoding CylC homolog(s).

***Desmonostoc muscorum* LEGE 12446**
**Cluster 2**

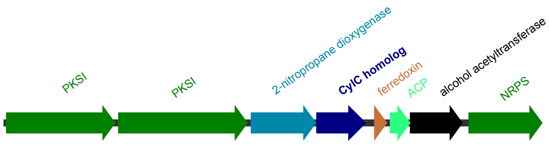


***Nostoc* sp. LEGE 12450**
**Cluster2**

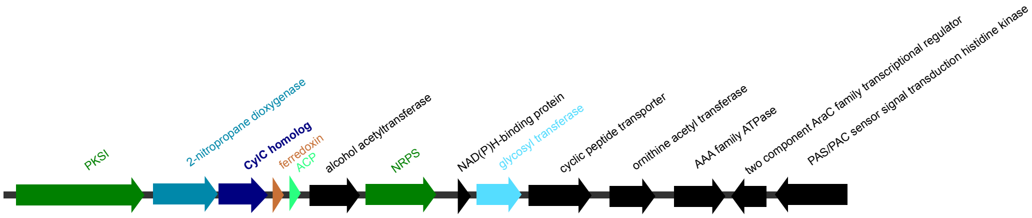


***Nostoc sp.* LEGE 12447**
**Cluster 2**

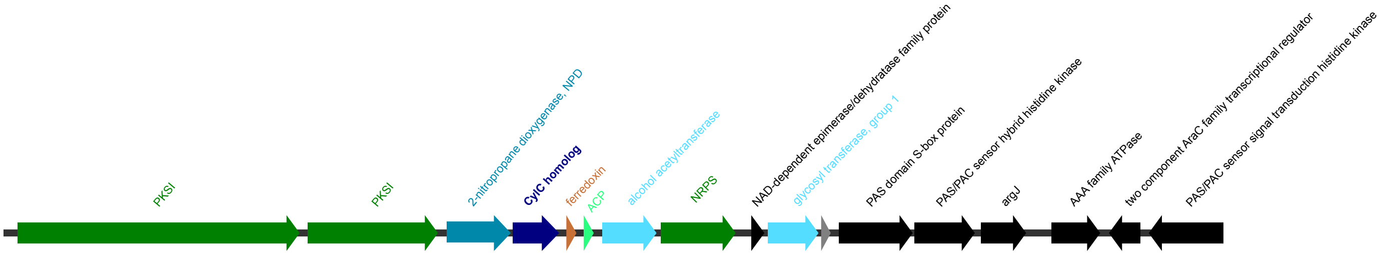


**Figure S10.** Nitronate monooxygenase-containing biosynthetic gene clusters encoding a CylC homolog.

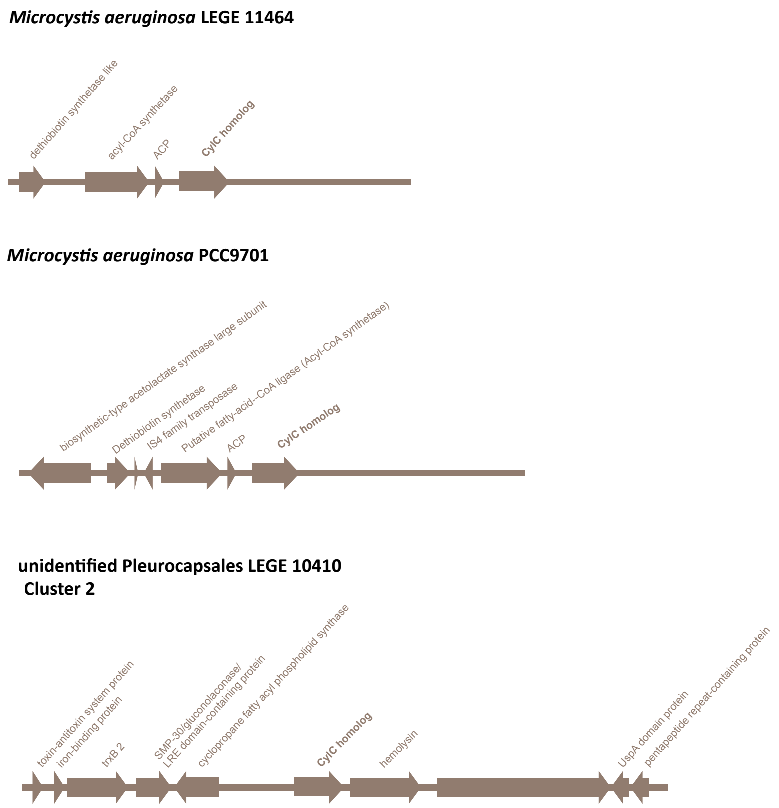


**Figure S11.** Unclassified (likely incomplete) biosynthetic gene clusters encoding a CylC homolog.

**Table S4.** BLAST search of Rieske-containing BGCs genes from *Calothrix brevissima* NIES 22 against *Synechocystis* sp. PCC 6803.

| Gene | Annotation | PCC 6803 ortholog | E-value | Identity | Query Coverage | Bit-Score |
|---|---|---|---|---|---|---|
| **NIES22_66770** | DUF962 domain-containing protein | none | | | | |
| **NIES22_66780** | Rieske aromatic ring-hydroxylating dioxygenase | sll1849 | 8.68E-97 | 68.6 | 61.63 | 311.227 |
| **NIES22_66790** | hypothetical | sll0263 | 2.85E-67 | 46.4 | 84.92 | 222.246 |
| **NIES22_66800** | Rieske aromatic ring-hydroxylating dioxygenase | sll0264 | 5.74E-137 | 60.2 | 97.01 | 426.402 |
| **NIES22_66810** | ferritin-like diiron protein | sll0265 | 8.33E-74 | 43.3 | 98.67 | 243.817 |
| **NIES22_66820** | hypothetical | slr0326 | 1.31E-17 | 67.1 | 78.65 | 72.789 |
| **NIES22_66830** | CylC-like dimetal-carboxylate halogenase | sll0266 | 5.63E-158 | 60.8 | 94.37 | 489.574 |
| **NIES22_66840** | hypothetical | none | | | | |
| **NIES22_66850** | ferredoxin | ssl2559 | 1.79E-28 | 53.4 | 93.62 | 107.071 |
| **NIES22_66860** | Rieske aromatic ring-hydroxylating dioxygenase | sll1297 | 3.15E-147 | 61.5 | 100 | 456.447 |
| **NIES22_66880** | sterol desaturase or sphingolipid hydroxylase | sll1510 | 1.88E-59 | 45.9 | 99.59 | 199.904 |
| **NIES22_66890** | acyl transferase | sll1511 | 3.08E-56 | 51 | 76.12 | 188.734 |

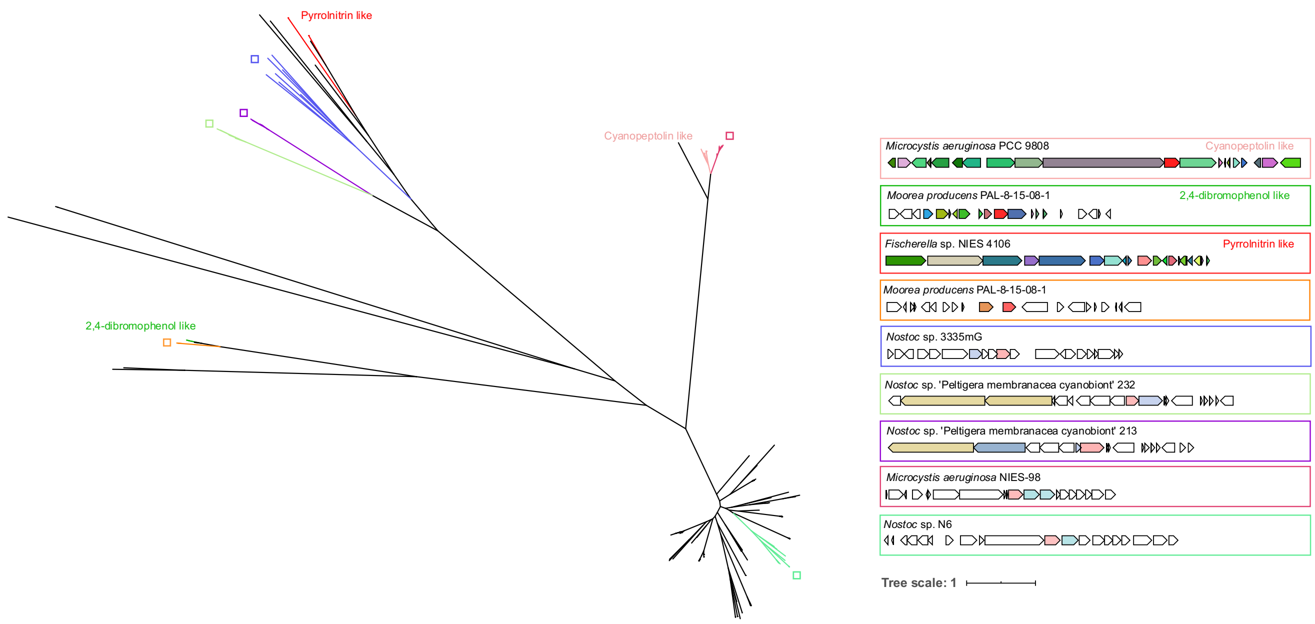


**Figure S12.** Phylogenetic tree of FAD-dependent halogenases based on CORASON outputs with illustrative BGC architectures.

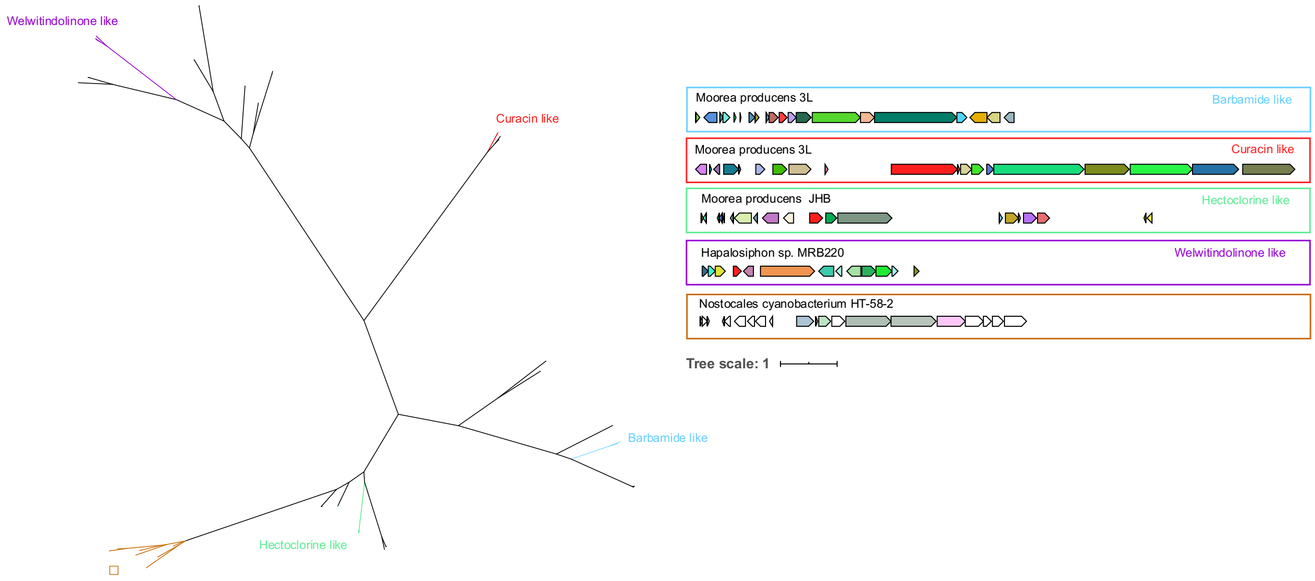


**Figure S13.** Phylogenetic tree of nonheme iron-dependent halogenases based on CORASON outputs with illustrative BGC architectures.

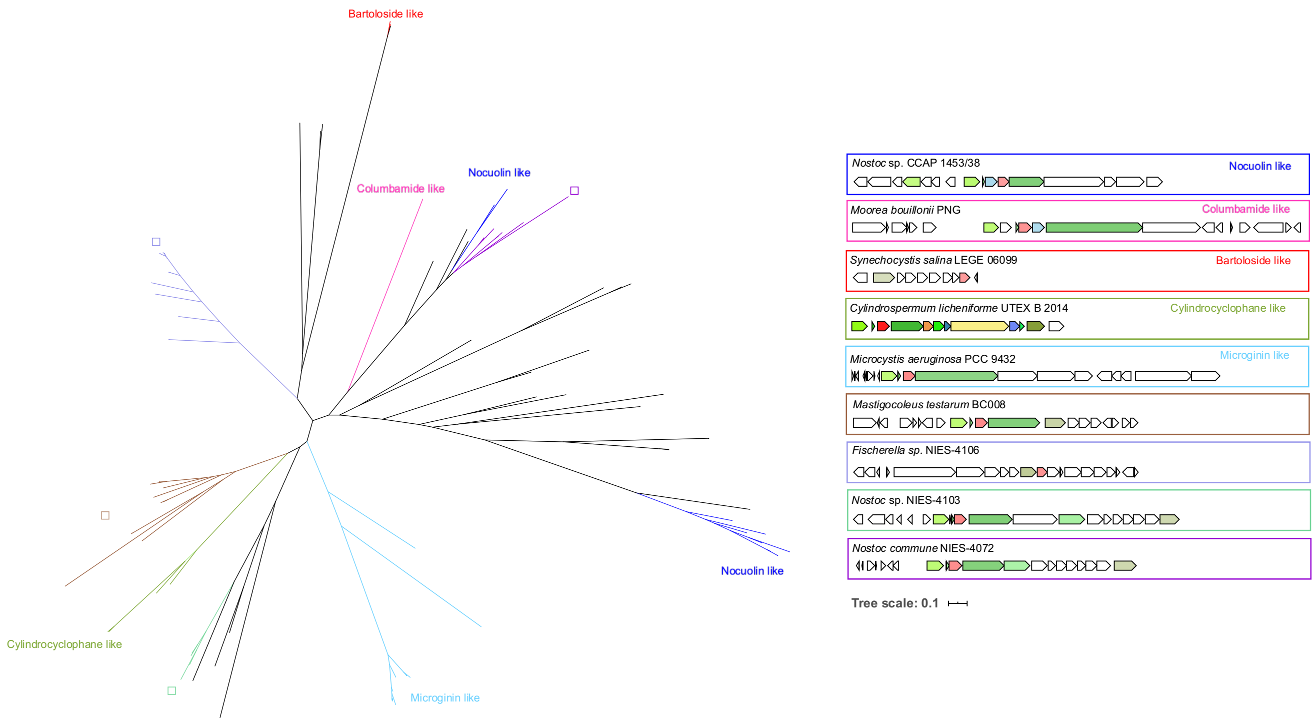


**Figure S14.** Phylogenetic tree of dimetal-carboxylate halogenases based on CORASON outputs with illustrative BGC architectures.
